# A Benchmarking Framework for Comparative Evaluation of Low-Complexity Region Detection Tools in the Human Proteome

**DOI:** 10.64898/2026.01.24.701293

**Authors:** Anirjit Chatterjee, Nagarjun Vijay

**Author notes:** Author for Correspondence: Anirjit Chatterjee, Nagarjun Vijay.

## Abstract

Low-complexity regions (LCRs) are compositionally biased segments of proteins that play critical roles in molecular recognition, structural flexibility, and phase separation. Yet, their accurate detection remains challenging due to methodological variability among computational tools. In this study, we conducted a comprehensive benchmarking of eight widely used LCR detection methods (with different parameter settings) across the *Homo sapiens* proteome. A modular computational framework was developed to systematically compare LCR characteristics, including residue-centric analyses such as length distributions and coverage percentages. Protein-centric analyses consisted of compositional bias, amino acid composition, and Shannon entropy. Consensus analyses revealed that regions detected by multiple tools were typically longer, more repetitive, and compositionally purer, suggesting stronger structural or functional relevance. Jaccard similarity matrices demonstrated distinct clustering patterns among algorithms based on shared detection principles. Additionally, entropy and purity analyses highlighted fundamental differences in sequence complexity captured by each tool. Together, these results provide a unified, reproducible framework for evaluating LCR detection performance and offer practical guidelines for reliably annotating low-complexity regions in proteome-scale studies.

## INTRODUCTION

In recent years, low-complexity regions (LCRs) have gained recognition as significant drivers of the genome. ^1–3^. While the precise definition of LCRs remains a topic of debate, they are broadly accepted as sequences characterised by repetitive amino acid patterns (e.g., ‘ATGGGGATTT’ or ‘LKSSSQQQQSSSK’). Historically referred to as Compositionally Biased Regions (CBRs) ^4^, Cryptic Repeats ^5^ and Tandem Repeats ^6^, these regions are now commonly termed LCRs ^7^. Defined by their reduced amino acid diversity, LCRs range from simple single amino acid repeats to stretches with limited residue types.^8–10^ Found in 18–20% of human proteins and is more prevalent in mammals^11,12^, LCRs serve as evolutionary "tuning knobs," fostering adaptation and diversity ^13^. At a more granular level, the evolution of low-complexity regions is often driven by nucleotide-level mechanisms like replication slippage and repeat instability. ^14^

Previously overlooked due to their intrinsically disordered nature and labelled as part of the “Dark Proteome”, LCRs are now recognised for their roles in phenotypic variation and disease mechanisms. For example, the polyQ-polyA repeat ratio in RUNX2 influences vertebrate facial morphology.^15^, while expanded polyQ tracts within LCRs contribute to neurodegenerative disorders such as Huntington’s disease (HD) and spinal and bulbar muscular atrophy (SBMA) ^16^. Functionally, LCRs are pivotal in biological processes like cell signalling^17^, RNA processing ^18^, transcription regulation ^13,19,20^, DNA repair, and neurogenesis^12^. Structurally, LCRs regulate protein solubility and folding, as seen in prion-like LCRs^21^, and play roles in RNA binding and phase separation through motifs like RGG/RG^22^. They can form labile cross-β polymers and hydrogel droplets.^23^, highlighting their versatility in molecular assembly and cellular organisation. In *Plasmodium falciparum*, longer proteins have been shown to have preferential enrichment of LCRs.^24^

To advance the study of LCRs, various detection methods have been developed in recent times, each employing unique algorithms. However, the diversity of these tools necessitates benchmarking to compare and analyse their performance.^25^ Previous efforts to evaluate and compare low-complexity region (LCR) detection methodologies have addressed multiple complementary aspects, including formalising operational definitions of LCRs,^26^ systematically reviewing tandem repeat detection algorithms,^27^ and developing integrative platforms such as PlaToLoCo—the first web-based meta-server that unifies outputs from diverse LCR detection methods to enable comparative visualisation and interpretation of protein low-complexity landscapes.^28^ Recently, a database called LCRAnnotationsDB has been developed to include unified functional and structural information from a large number of publicly available protein databases.^29^ This study evaluates eight LCR detection methods—fLPS, fLPS 2.0, AlcoR, Dotplot, SEG, LCRfinder, T-REKS and XSTREAM —selected for their availability as command-line tools and representation of diverse detection approaches. Previously, SEG was widely used for masking LCRs in BLAST searches ^30^, while at present, fLPS and its updated version, fLPS 2.0, focus on identifying compositionally biased regions ^31,32^. AlcoR employs bidirectional compression for mapping and visualising LCRs ^33^, LCRfinder examines LCRs from an evolutionary perspective^34^, and Dotplot leverages dimensionality reduction to analyse LCR sequence space ^10^. T-REKS and XSTREAM both use algorithms to detect the presence of Tandem Repeats within sequences ^35,36^.

This study analyses 20,610 sequences across the *Homo sapiens* proteome^10^ using these methods, offering insights into their detection capabilities. Various parameters were taken into account to analyse the differences and similarities between the detections by these tools. Studies were also made to look into the consensus of the results of the detections. Efforts were also made to define the boundaries of the Low Complexity space,^26^ using disordered regions as background and taking into account sequences detected by the method consensus. Additionally, plots of sequence complexity landscapes were analysed across tools to study the variations in complexity. Evaluation metrics were utilised to study the performance of the methods against various parameters. This research represents a comprehensive benchmarking effort in the field, aiming to encourage the development of more robust studies and methodologies for LCR analysis.

## METHODOLOGY

### Selection of LCR Detection Tools for Benchmarking

A diverse set of computational tools was selected to comprehensively benchmark the detection of low complexity regions (LCRs) in protein sequences. The tools were grouped into three broad categories based on their underlying methodologies and usage in the literature. The first category, popular tools, included SEG and fLPS (along with its updated version, fLPS 2.0), which are widely used in LCR studies due to their established reliability and computational efficiency. The second category comprises tools designed specifically for tandem repeat detection, including XSTREAM and T-REKS, which identify repetitive motifs that often characterise certain types of LCRs. The third group encompassed novel and emerging methods, such as a custom Dotplot-based approach, LCRFinder, and AlcoR. These tools were chosen for their innovative algorithms and potential to provide complementary insights. AlcoR, in particular, utilises alignment-free techniques for LCR simulation, mapping, and visualisation. Tool selection was informed by recent literature, including methodologies described in Lee et al.^10^ and Silva et al.^33^, ensuring the inclusion of both traditional and state-of-the-art approaches for robust and comparative analysis.

### Tools used

SEG is a computational tool developed by Wootton and Federhen (1993) ^30^ to identify low-complexity regions (LCRs) in amino acid sequences. It operates by calculating the local sequence complexity within a sliding window using an information-theoretic measure of sequence entropy. Regions with lower-than-expected complexity—where a limited set of residues recur frequently—are masked or flagged as low complexity. This approach helps distinguish compositionally biased or repetitive regions from more diverse segments of the sequence. SEG is widely used in sequence analysis pipelines, especially in database searches, to reduce spurious alignments caused by non-informative, repetitive regions.

fLPS (fast Lowest Probability Subsequence) is a computational tool designed to detect compositionally biased or low-complexity regions (LCRs) in protein sequences with high speed and statistical rigour ^31^. It identifies segments where the amino acid composition deviates significantly from what would be expected under a random model, using a probabilistic framework to estimate the likelihood of observing such biases. Unlike earlier tools such as SEG, fLPS emphasises statistical significance rather than purely entropy-based measures, allowing for more sensitive and precise detection of compositional biases across diverse proteomes.

fLPS 2.0 is an improved version that enhances both the accuracy and efficiency of LCR detection ^32^. It introduces refined background models and better handling of mixed or weakly biased regions, enabling more nuanced identification of low-complexity segments. The updated algorithm also supports larger datasets and provides greater flexibility in parameter tuning, making it suitable for large-scale proteome analyses and comparative studies of sequence bias.

AlcoR (Algorithm for Low-Complexity Region detection) is a computational tool developed to identify low-complexity and compositionally biased regions in protein sequences through a statistically grounded, alignment-free approach ^33^. It quantifies amino acid repetitiveness and local compositional bias by analysing the recurrence and arrangement of short sequence motifs. Unlike earlier methods such as SEG or fLPS, AlcoR integrates both compositional and positional information, allowing it to detect a wider range of repetitive or biased regions, including subtle or irregular patterns. The algorithm is optimised for large-scale proteome analyses, offering high speed, adjustable sensitivity, and compatibility with downstream characterisation of detected low-complexity regions.

T-REKS (Tandem Repeats Extraction and Analysis Tool) is a computational method designed to identify tandemly repeated motifs within protein sequences ^35^. It detects repeats by clustering identical or highly similar short subsequences (k-mers) and refining their alignment to define repeat boundaries and consensus motifs. T-REKS combines sequence similarity searches with statistical evaluation to distinguish true tandem repeats from random similarities, allowing accurate detection of both perfect and imperfect repeats. The tool is particularly effective in analysing proteins with modular or repetitive architectures, such as surface and structural proteins, identifies short tandem repeats which are part of low-complexity domains, and is frequently used in large-scale proteome analyses to study sequence repetitiveness and evolutionary divergence.

XSTREAM (eXtendedSTRing Repeat detection program) is a versatile algorithm developed to detect and characterise repetitive sequence motifs in proteins and nucleic acids ^36^. It identifies both tandem and interspersed repeats, accommodating imperfect, variable-length, and nested repeat structures. XSTREAM uses pattern matching combined with flexible scoring and alignment parameters to recognise repeats that may evolve through mutation, insertion, or deletion. The tool reports detailed information about each repeat family, including consensus motifs, copy number, and sequence divergence. Its strength lies in its ability to detect complex and degenerate repeat architectures that are often missed by simpler repeat-finding algorithms, making it valuable for studying protein domains, low-complexity regions, and the evolutionary dynamics of repetitive sequences.

Dotplot analysis is a graphical method used to visualise sequence self-similarity and repetitive patterns within proteins or nucleic acids. In this approach, a sequence is compared against itself (self-dotplot) or another sequence by plotting matching residues as dots on a two-dimensional grid—one axis representing the query sequence and the other the target ^10^. Continuous diagonal lines indicate regions of similarity or internal repeats, while scattered dots suggest randomness or compositional variation. When applied to proteins, dotplots effectively reveal low-complexity regions (LCRs), tandem repeats, and internal duplications that might not be easily detected by linear algorithms. This method provides an intuitive, alignment-free visualisation of repetitive organisation and has become a fundamental step in studying sequence complexity and structural modularity across proteomes.

LCRFinder is a computational method developed to identify low-complexity regions (LCRs) in protein sequences based on local amino acid composition bias and sequence repetitiveness. ^34^ It employs a statistical framework that scans protein sequences to detect short segments where specific amino acids or motifs occur more frequently than expected under a random distribution. Unlike earlier tools that rely primarily on entropy or window-based masking, LCRFinder integrates compositional enrichment analysis with pattern clustering, allowing it to sensitively detect both simple and complex forms of low complexity.

### Selection of the Dataset

For this study, the human proteome was obtained from the UniProt database, ensuring a high-quality and comprehensive dataset for downstream analyses. A multi-protein FASTA file containing 20,610 protein sequences was selected. To maintain dataset integrity and biological relevance, only one representative protein per gene was retained, excluding all isoforms. This rigorous filtering ensured minimal redundancy and high annotation quality, providing a robust foundation for the analysis of low complexity regions (LCRs). The dataset selection strategy follows the guidelines established by Lee et al. ^10^.

### Planning the Methodology

The proteome was analysed for LCR detection by the selected tools. Each tool gave outputs in separate formats. To ensure a uniform framework for benchmarking diverse LCR detection tools, all tool outputs were standardised into a common BED-like format, using custom Python scripts. This modification allowed direct comparison of predicted low-complexity regions (LCRs) across tools, independent of their native output formats. Each BED entry captured the essential parameters: protein identifier, start and end positions of the detected LCR.

### Data Aggregation, Processing, and Visualisation

To systematically evaluate and compare the performance of multiple low-complexity region (LCR) detection tools, categorised output files were collected across five major analytical dimensions: LCR length distribution, LCR coverage percentage, LCR counts per protein, amino acid composition, and Shannon entropy diversity. Each dataset was generated using distinct LCR detection tools and operating modes and stored as tab-separated values (TSV) files.

Custom Python scripts (in GitHub repository) were used to process the raw outputs from each tool. These scripts calculated LCR lengths from start and end coordinates, determined the percentage of each protein sequence covered by LCRs, quantified amino acid compositions, and computed Shannon entropy values to evaluate compositional diversity. For comparative analyses, LCRs were assigned to fixed length bins (0–10, 10–20, 20–50, 50–100, 100–200, and >200 amino acids), while LCR coverage percentages were categorized into 0–20%, 20–40%, 40–60%, 60–80%, and 80–100% bins. LCR counts per protein were grouped into 0, 1–5, 5–10, 10–15, and 15+ categories. Amino acid proportions were normalised to total residue counts, and entropy distributions were computed to assess the complexity of the detected regions.

All categorised datasets were subsequently processed and visualised in R (version ≥4.0.0)^37^ using the tidyverse^38^, ggplot2^39^, viridis^40^, reshape2^41^, dplyr^42^, and tidyr^43^ Packages. Data from all analyses were harmonised by extracting tool identifiers from filenames using regular expressions, merging dataframes iteratively, and converting categorical variables into ordered factors to ensure consistent axis ordering across plots.

Visualisation was performed using ggplot2 and complementary packages including ggpattern, Polychrome, and patchwork. Heatmaps were generated to display LCR length and coverage distributions, where colour gradients represented frequency intensities. Stacked bar plots illustrated LCR counts per protein and amino acid compositions across tools, while boxplots summarised the Shannon entropy distributions. Additional composite visualisations integrated motif distribution, entropy variation, and sequence purity metrics, enabling a comprehensive, multi-dimensional comparison of the sensitivity, selectivity, and compositional characteristics of the evaluated LCR detection methods.

### Analysis of Common LCR Properties Across Detection Tools

To characterise the shared properties of low-complexity regions (LCRs) identified by different computational tools, a comparative overlap analysis was performed using BEDTools (v2.30.0)^44^. The *bedtools multiinter* command was employed to identify genomic coordinates of common LCRs detected across multiple methods, followed by stratification of these intersected regions based on the number of tools detecting them (ranging from 1 to 13). Using the *bedtools getfasta* command, amino acid sequences corresponding to each intersected region were extracted from the *Homo sapiens* reference proteome FASTA file for downstream compositional and motif analyses.

Custom Python scripts were used to determine the most frequently occurring peptide motifs within these shared regions. Each region was annotated with its dominant motif and classified into one of four categories—Monopeptide, Dipeptide, Tripeptide, or Other—based on regular expression–based pattern matching. The top twenty most abundant peptide repeats were summarised, and their proportions were visualised in R (version ≥4.0.0) using a custom graphical workflow implemented with the ggplot2, ggpattern, dplyr, Polychrome^45^, and patchwork^46^ Packages.

The visualisation pipeline consisted of three main components:

1. Motif Composition Plot – A stacked bar plot displaying the relative abundance of dominant peptide motifs across regions detected by varying numbers of tools. Distinct fill patterns (solid, stripe, crosshatch) were used to differentiate monopeptides, dipeptides, and tripeptides, respectively.
2. Entropy Analysis – Shannon’s Diversity Index was calculated for each region to quantify compositional heterogeneity. Boxplots were generated using color-blind–friendly palettes to depict entropy distributions across consensus levels, with outliers excluded for clarity.
3. Purity Analysis – Simpson’s Purity Index was computed to measure the dominance of single amino acids within regions. Corresponding boxplots were produced in parallel with the entropy distributions for direct comparison.

All three visual components were combined into a composite multi-panel figure using the patchwork package, enabling simultaneous interpretation of motif proportions, entropy variability, and compositional purity across increasing tool consensus levels.

Together, these analyses provided a unified view of how increasing agreement among LCR detection tools correlates with compositional simplicity, reduced entropy, and enrichment for highly repetitive peptide motifs, thus revealing the convergent tendencies of algorithms toward strongly biased low-complexity regions.

### Visualisation of low-complexity (LC) diagrams

Low-complexity (LC) regions were visualised in a two-dimensional sequence-complexity space defined by mutation percentage and the fractional contribution of the most frequent amino acid. LC regions identified at different consensus levels (k = number of methods) were aggregated across output files following numeric validation and removal of incomplete entries. Kernel density estimation was applied to generate two-dimensional contour representations of LC distributions, allowing direct comparison of density patterns across consensus tiers. Contours were coloured by consensus level, with line widths scaled to reflect tier order. All density plots were rendered using fixed 0–100% axes to ensure consistent scaling across methods. (plot1.r)

To quantify shifts in central tendency and dispersion across consensus tiers, median mutation percentage and compositional dominance were computed for each tier, together with interquartile ranges (IQRs). These summary statistics were visualised as centroid points with orthogonal IQR crossbars. Centroids corresponding to successive consensus tiers were connected by directional arrows to illustrate systematic transitions in LC composition with increasing consensus stringency. This representation captures both variability and directional trends while remaining robust to outliers. (plot2.r)

To assess cumulative occupancy of LC space, mutation percentage and compositional dominance values were discretised into fixed 2% × 2% bins. For each bin, the highest consensus tier observed among LC regions falling within that bin was recorded. This information was visualised as a tile-based heatmap, where colour intensity reflects the maximum level of cross-method support present in each region of sequence-complexity space. Bin geometry, coordinate limits, and aspect ratios were held constant to preserve spatial interpretability. (plot3.r)

Finally, tier-specific LC density patterns were examined by generating independent binned density maps for each consensus tier using identical bin sizes and axis limits. Within each tier, the number of LC regions per bin was counted and visualised as faceted heatmaps. A square-root transformation was applied to bin counts to stabilise variance and improve visual contrast between sparse and dense regions. This small-multiple approach enables direct comparison of how LC landscape occupancy changes as consensus stringency increases. (plot4.r)

### Visualisation of Entity Retention Across Purity Levels

To assess and compare the performance of multiple computational methods in retaining entities across varying purity thresholds, a custom workflow was implemented in R (version ≥4.0.0) using the ggplot2, dplyr, tidyr, and Polychrome packages. Tab-delimited data containing the proportion of entities retained by each method at different purity levels were imported and grouped by detection method.

For each method, line plots modelled the relationship between purity level and entity retention.

A distinct colour palette was generated using the Polychrome package to ensure clear visual distinction among methods. The processed data were visualised using ggplot2 as a line plot, with purity level on the x-axis and the proportion of entities retained on the y-axis. Each method was represented by a unique colour-coded line and corresponding points.

Consistent aesthetic parameters— including uniform axis formatting, font sizes, and legend scaling—were applied to maintain clarity and visual harmony across figures.

### Jaccard Similarity and Overlap Analysis Between LCR Detection Tools

To evaluate and compare the degree of overlap and similarity among various computational methods for low-complexity region (LCR) detection, a correlation-style heatmap was constructed using custom R scripts. All analyses were conducted in R (version ≥4.0.0) with the corrplot^47^, tidyverse and reshape2 packages for data manipulation and visualisation.

A Jaccard similarity matrix was derived from actual LCR coordinate overlaps encompassing eight methods, including AlcoR (Mode 1 and Mode 2), SEG (default, intermediate, and strict modes), fLPS (default and strict), fLPS 2.0 (default and strict), Dotplot, LCRFinder, T-REKS, and XSTREAM. Pairwise Jaccard coefficients were computed using the *bedtools jaccard* command, and the resulting similarity file (*human_jaccard.tsv*) was imported into R, formatted as a numeric matrix, and harmonised with the overlap dataset.

A figure was then produced which depicted the Jaccard similarity matrix with a diverging white–red colour scheme to represent low-to-high similarity levels.

### Estimating the boundaries of low-complexity (LC) sequence space

Low complexity sequence space has been defined with transient boundaries,^26^ but is not defined by a single physical rule but emerges from reduced sequence diversity, compositional bias, and tolerance to mutation. To ground this abstract concept biologically, we framed LC space using experimentally supported disorder as a reference. We used DisProt^48^ annotations, which represent experimentally validated intrinsically disordered regions, and missing residues in PDB structures, which indicate regions that fail to adopt stable conformations in structural experiments. These datasets capture functional and structural manifestations of disorder, respectively, and provide biologically meaningful anchors for interpreting low-complexity sequence patterns. Such disordered regions served as background. Each sequence segment was represented using two intuitive features: mutation percentage, which reflects how freely a region can change without disrupting its role, and percentage of the most frequent amino acid, which reflects how compositionally biased the sequence is. Together, these features summarise how information-rich or information-poor a sequence region is.

In the first approach, the empirical LC boundary was estimated using binning (*plot_LC_boundary_1.r*). LC space was discretised into a regular two-dimensional grid defined over mutation percentage (x-axis) and amino-acid dominance (y-axis), using bins of width Δ = 2%. Each sequence segment with coordinates (*x,y*) was assigned to a bin

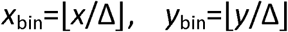

For each bin *(i,j)*, we counted the number of LC segments *n_ij_^+^*and background segments *n_ij_^−^*. The probability that a segment in that bin belongs to an LC region was estimated using a smoothed frequency ratio:

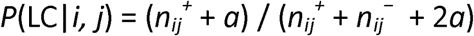

with *a*=1. This small correction prevents unstable estimates in sparsely populated bins and reflects prior uncertainty rather than assuming zero probability.

To evaluate how well these probabilities separate LC from non-LC segments, we performed receiver operating characteristic (ROC) analysis on held-out data. For a probability threshold t, predictions were defined as LC if *P ≥ t*. The true-positive rate (TPR) and false-positive rate (FPR) were computed as

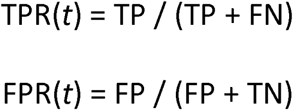

The optimal threshold *t\** was selected by maximising Youden’s J statistic,

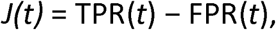

which balances sensitivity and specificity. The LC boundary was then defined as the contour in feature space where *P*(LC)=*t*^∗^. This contour represents a confidence boundary, not a strict biological delimiter.

The second approach estimates LC boundaries using a smooth probabilistic surface rather than discrete bins using a generalised additive model (GAM) (*plot_LC_boundary_2.r*). We modelled the probability that a segment belongs to an LC region as a continuous function of mutation percentage *x* and amino-acid dominance *y*:

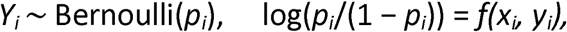

where *Y_i_*indicates LC status and *f(x,y)* is a smooth two-dimensional function learned from the data.

This function was represented using penalised regression splines, which allow the surface to adapt to data while discouraging unnecessary complexity. Smoothness penalties ensure that the inferred LC probability varies gradually across sequence space, consistent with biological expectations of gradual transitions rather than sharp cutoffs.

Predicted probabilities *p̂_i_* were evaluated using ROC analysis as above, and an optimal threshold *t*^∗^ was selected via Youden’s J statistic. To visualise the LC boundary, probabilities were evaluated over a regular grid covering the domain [0,100]×[0,100], yielding a posterior probability surface

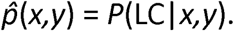

The LC boundary was extracted as the contour where *p̂*(*x,y*) = *t\**.

Together, these complementary approaches show that LC regions do not occupy a sharply defined territory in sequence space. Instead, LC space emerges as a probability envelope, shaped by entropy reduction, compositional bias, and biological disorder constraints. DisProt annotations and missing PDB residues provide biological grounding, while probabilistic modelling reveals that confidence in low complexity increases gradually rather than abruptly. The explicit mathematical formulation underscores that LC boundaries are inherently fuzzy and context-dependent, reflecting a continuum of sequence information rather than a binary state.

### Visualisation of the most frequent amino acid and mutation percentage

To explore the interplay between amino acid composition, periodicity, and structure in protein regions, a two-dimensional (2D) diagram plotting compositional bias (y-axis) versus repetitiveness (x-axis) was constructed, according to the model presented by Lee et al. 2022^26^. The analysis has focused on sequence regions of 10–50 residues, a length optimal for balancing statistical robustness and biological relevance. For each region, two complexity metrics were computed: (1) the frequency of the most abundant amino acid as a proxy for compositional bias, and (2) the proportion of residues that must be mutated to convert the region into a perfect repeat, representing repetitiveness. Sample sequences were analysed to demonstrate these relationships, such as ACDEFEGEIE, where the highest frequency amino acid (E, 40%) sets y, and periodicity is estimated by calculating minimal mutational distance to a perfect repeat (e.g., EEEEEEEEEE, 60%). The complexity triangle represents a continuous sequence landscape defined by amino acid compositional dominance and mutational distance to perfect repetition. Sequences with uniform amino acid usage (e.g., globular proteins) cluster at the lower-right of the diagram, while perfect and homorepeat sequences occupy the left side. Intermediate sequences (e.g., direpeats, tripeats) span between these extremes.

Utilising this model, we mapped selected proteins with low-complexity regions (LCRs) and unbiased regions into the diagram to visualise how their sequence characteristics relate to their potential structural tendencies and LCR classification.

The two-dimensional diagram was broadly divided into 3 segments:

1. LCR segment (X < 50 and Y > 50)
2. CBR segment (X < 50 and Y < 50)
3. HCR segment (X > 50 and Y < 50)

The exact boundaries between regions of various complexity are difficult and ambiguous to identify (as attempted in the above subsection). For operational classification, this space was subdivided into low-complexity (LCR), compositionally biased (CBR), and high-complexity (HCR) regions using 50% thresholds along both axes. These thresholds were chosen for interpretability rather than as strict biological cutoffs. A dominance value of 50% separates sequences primarily governed by a small subset of residues from those with more evenly distributed amino acid usage, consistent with classical entropy- and composition-based definitions of low complexity.^49^ Similarly, a mutation requirement of 50% distinguishes sequences that are close to repetitive or biased patterns from those requiring extensive substitutions to reach such states, aligning with prior treatments of repeat degeneracy and compositional bias.^50,51^ It is to be emphasised that these boundaries are heuristic and intended for comparative contextualisation rather than strict biological classification.

A Python-based computational pipeline was developed to evaluate the compositional complexity of low-complexity regions (LCRs) in protein sequences and to detect them using a sliding-window approach. The workflow accepts a protein FASTA file and a corresponding BED file as input to extract LCR coordinates, employing a sequence window size of 20 amino acids with a sliding window of 10 amino acids (using *seq_parser.py*). For each extracted region, Shannon entropy was calculated to quantify sequence diversity, while amino acid frequencies were analysed to identify the most abundant residue and its relative contribution. Repetitive sequence motifs (k-mers) of varying lengths were detected, and the minimal mutation percentage required to achieve perfect periodicity was estimated. The resulting LCRs were visualised as density plots with repetitiveness and compositional bias represented on the x- and y-axes, respectively (using *complexity_plot.py*), and all computed parameters—including entropy, dominant residue, k-mer composition, and mutation metrics—were compiled into a structured tab-delimited output file.

### Annotating “Positives” and “Negatives”

To quantify the agreement between low-complexity regions (LCRs) detected by different methods, we computed true positives (TP), false positives (FP), false negatives (FN), and true negatives (TN) using a residue-level comparison framework. Reference LCRs were obtained from the resulting sequences generated from the pipeline in the above subsection (only those regions which had Mutation% % < 50 and Most Frequent Amino Acid% % > 50 were annotated as LCRs), whereas predicted LCRs were taken from the result bed files of the various methods used.

All protein sequences were treated as linear sequences of residues. Each residue was assigned a binary label indicating whether it belonged to an LCR according to the prediction set, the reference set, or both. A residue was considered “positive” if it fell within an annotated LCR interval and “negative” otherwise. A custom bash script (*conf_mat.sh*) was written to perform this comparison.

True positives were defined as residues demarcated as LCRs that also overlapped with each method-predicted LCR. False positives corresponded to residues demarcated as LCRs by the sliding-window method but not predicted as LCRs by the methods. False negatives were residues predicted as LCRs by the methods that were not recovered by the sliding-window–based segmentation approach. True negatives comprised residues not predicted as LCRs in either dataset. Counts were aggregated across all proteins prior to metric calculation.

### Visualisation of Tool Performance Across Multiple LCR-Associated Parameters

To comprehensively evaluate the performance of various low-complexity region (LCR) detection tools across multiple quantitative biological parameters, we developed a modular visualisation pipeline in R (v4.0.0 or later). The script incorporated the ggplot2, dplyr, tidyr, cowplot^52^, patchwork, and readr^53^ libraries to generate comparative line plots for both true positive rate (TPR) and false positive rate (FPR) across distinct feature categories, which were defined as:

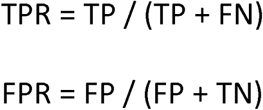

Four key LCR-related parameters were analysed: gene length, the number of LCRs per gene, the percentage of gene length covered by LCRs (LCR coverage), and the entropy ratio between LCRs and entire genes. For each parameter, precomputed TPR and FPR values for 8 different computational tools (with 13 parameter settings) were loaded from tab-delimited files and visualised using consistent formatting.

Gene length categories (1–10), LCR number (1-6), and LCR coverage per cent bins (5%, 10%, 15%, 20%, >20%) were treated as ordered categorical variables. Similarly, entropy ratios (0.2–1.0) were modelled as ordered factors. All plots employed a shared colour palette to maintain consistent tool representation. A common minimalist theme was defined and applied across all figures to ensure visual uniformity and clarity.

For each parameter, two line plots were generated—one for TPR and one for FPR—where tool performance was plotted against the relevant parameter category. The eight individual plots were grouped into four panels (TPR and FPR for each parameter) and arranged in a 2×2 grid using the patchwork and cowplot packages.

### Quantitative Comparison of LCR Detection Tools Based on Mean TPR and FPR Across Multiple Parameters

To summarise the overall performance of 13 computational tools in detecting low-complexity regions (LCRs), we calculated and visualised the mean true positive rate (TPR) and false positive rate (FPR) under four different biological stratifications: gene length, number of LCRs per gene, LCR coverage percentage, and entropy ratio between LCR and gene.

All analyses and visualisations were performed in R (v4.0.0 or later) using the packages ggplot2, dplyr, tidyr, and patchwork. Preprocessed tab-delimited data files were read from the respective directories for each stratification. Within each dataset, performance metrics (TPR and FPR) were grouped by tool, and the mean values were calculated.

Each parameter yielded a pair of grouped bars per tool—one for mean TPR and one for mean FPR—to visually separate the metrics. The tools were ordered along the x-axis by descending value for each plot to enhance interpretability. Custom axis themes and font sizes were applied through additional formatting.

### Quantification and visualisation of structurally disordered regions

To perform the validation of structurally disordered regions in the Low Complexity sequence space, the entire dataset of experimentally verified disordered regions was selected from DisProt ^48^ and those regions corresponding to *Homo sapiens* were filtered out (along with the coordinates of the disordered regions) into a bed file. These were further classified based on the experimental methods used to study the disordered regions. Based on the complexity triangle, each region was plotted into the Low Complexity sequence space, with each method being represented separately by distinctly coloured points.

Furthermore, protein regions lacking atomic coordinates in experimentally determined structures were used as structural evidence consistent with conformational disorder. Missing-residue annotations were obtained from UniProt–PDB residue mapping files by identifying residues present in the UniProt canonical sequence but absent from corresponding PDB atomic coordinates. Missing residues were interpreted as conservative structural indicators compatible with disorder, acknowledging alternative explanations such as experimental limitations, crystal packing effects, or transient structural states. Similar to the previous method, these segments were plotted into the Low Complexity sequence space as a heatmap.

## RESULTS

### LCR detection methods are different from each other

#### Variation in LCR Length Distribution

**Fig. 1A** demonstrates unique detection patterns influenced by the design of each tool. This analysis is residue-centric, focusing on the length distribution of individual low-complexity segments irrespective of their protein context. The fLPS clan (fLPS, fLPS strict, fLPS 2.0, fLPS 2.0 strict) detects the largest number of LCRs, particularly in larger categories (50–200+ amino acids), with over 60,000 counts in the 200+ category. This is consistent with fLPS’s emphasis on detecting large compositional biases. Conversely, AlcoR (Mode 1 and Mode 2) is less selective, focusing on LCRs of lengths 10–50 residues, as expected with its statistical bias toward local significance. Dotplot also focuses on shorter LCRs, but prefers 10–50 residue regions associated with self-similarity over extended bias. SEG family (SEG, SEG intermediate, SEG strict) is biased toward finding extremely short regions (0–20 residues), in accordance with its entropy-based methodology for masking very uniform segments. LCRFinder takes this even further, with the majority of ultra-short motifs less than 10 residues. In contrast, T-REKS and XSTREAM, designed for periodic tandem repeats, identify fewer LCRs, primarily between 10–100 residues. In general, the distribution profiles demonstrate how algorithmic variation affects detection.

**Figure 1.**
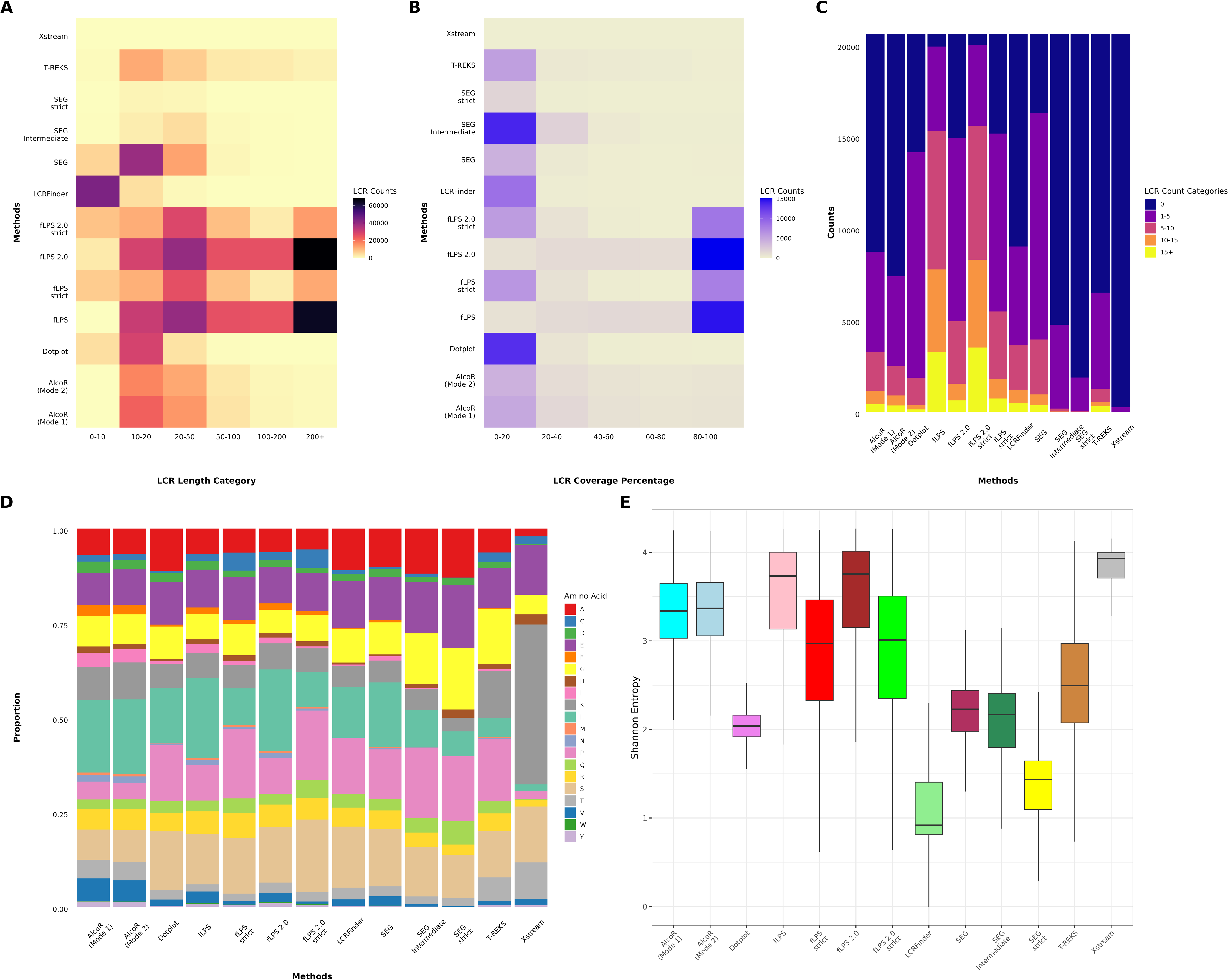
Comparative overview of Low Complexity Region (LCR) detection across multiple methods. **(A)** shows the total number of LCRs detected by each method across the analysed proteomes, where the x-axis lists the methods and the y-axis indicates the total LCR count. **(B)** represents the average LCR coverage per proteome, expressed as the percentage of total amino acids falling within LCRs. **(C)** compares the mean length of LCRs detected by each method, while **(D)** depicts the mean number of LCRs per protein sequence. Boxplots indicate variability across species, and horizontal lines mark median values. Together, these panels summarise the overall detection capacity, compositional coverage, and granularity of each computational approach.

### Variation in LCR Coverage

**Fig. 1B** reveals clear patterns among various LCR detection tools based on the fraction of residues within a protein annotated as low-complexity, providing a residue-centric view normalised to protein length. The fLPS set of tools (fLPS, fLPS strict, fLPS 2.0, and fLPS 2.0 strict) strongly identifies numerous proteins with 80–100% LCR coverage, indicating that these algorithms are very sensitive to long compositional biases and tend to mark large or whole proteins as low-complexity. Whereas programs such as AlcoR (Mode 1 and Mode 2) and Dotplot primarily find LCRs spanning merely 0–40% of protein sequences, the reason is the conservative nature that targets local high-scoring, low-complexity regions more than extended domains. The SEG family demonstrates mixed behaviour: SEG and SEG strict preferentially identify short LCRs (0–20% coverage), whereas SEG intermediate identifies a larger percentage of proteins in the 20–40% range as a result of its loose entropy threshold. In the same manner, LCRFinder prefers finding LCRs that span less than 40% of the protein, in accordance with its emphasis on small, repetitive motifs. T-REKS and XSTREAM also prefer to identify structured repeats, primarily in the 0–20% coverage bin. Tools either prefer broad coverage, such as fLPS, or localised identification, such as SEG, AlcoR, and Dotplot.

### Variation in LCR Counts

To complement residue-level analyses, we next examined LCR counts per gene, a protein-centric metric that captures how frequently low-complexity regions occur within individual proteins, independent of their total length or coverage. The outcomes were highly variable among methods, as seen in **Fig. 1C**. Some tools, like fLPS and fLPS 2.0, including their “strict” variants, found numerous genes with very high LCR counts, most notably in the 10–15 and 15+ bins. This indicates high sensitivity toward LCR-abundant genes but simultaneously brings the prospect of higher false positives. Dotplot exhibited modest capacity to detect multiple LCR-bearing genes, representing high detection and probably higher specificity than the fLPS-based approaches. SEG and SEG intermediate identified fewer numbers of highly LCR-abundant genes, rather showing well-distributed cases among the 1–5 and 5–10 ranges, representing a more cautious and sensitive detection pattern. SEG strict was more restrictive, with fewer LCR detections in total. LCRFinder and T-REKS had intermediate sensitivity, primarily detecting genes with 1–5 LCRs. AlcoR (both modes) and XSTREAM were the most conservative, with the majority of genes being assigned to "0 LCRs" or "1–5 LCRs," suggesting low sensitivity. Overall, these results align with previous findings: very sensitive methods such as fLPS variants identify multiple LCRs per gene, whereas Dotplot, SEG, AlcoR, and XSTREAM emphasise specificity.

### Amino Acid Proportion Variability

To determine the amino acid composition biases intrinsic to various LCR detection approaches, we compared the relative amino acid compositions of predicted LCRs from all tools, as seen in **Fig. 1D**. Extensive compositional profile differences were noted, mirroring each approach’s inherent detection strategy and selectivity. AlcoR (both Mode 1 and Mode 2) and XSTREAM identified LCRs most heavily enriched in a limited set of amino acids, specifically lysine (K) and leucine (L), showing an apparently strong preference for highly biased homopolymeric or oligomeric areas. Likewise, SEG strict also had a highly discriminatory profile, with LCRs most heavily dominated by leucine (L) and isoleucine (I) in line with its highly stringent entropy thresholds aimed at isolating the most extreme compositional simplicity areas. Conversely, fLPS, fLPS strict, and the fLPS 2.0 variants presented wider distributions of amino acids, which included a greater diversity of residues like glycine (G), serine (S), and proline (P). The trend indicates an ability to identify more diverse LCR types, including flexible and structurally disordered regions. Dotplot and SEG (intermediate and default mode) also featured relatively balanced composition, though being moderately enriched with small, compact residues such as G, S, and P, consistent with their sensitivity for disorder-prone, composition-biased motifs. LCRFinder and T-REKS exhibited an intermediate profile, preferring a lesser, though more heterogeneous, range of amino acids than the heavily selective approaches. In general, these findings suggest that although very strict tools like AlcoR and XSTREAM are very good at identifying sharply biased, straightforward LCRs, more general-purpose tools like fLPS variants and Dotplot identify a greater variety of LCR types.

### Shannon Entropy Variation

**Fig. 1E** shows the compositional complexity of the putative low-complexity regions (LCRs) by comparing their Shannon entropy distributions. For the standard 20–amino-acid alphabet, the theoretical maximum Shannon entropy is log₂(20) ≈ 4.32, corresponding to uniform amino acid usage; lower values indicate increasing compositional bias. Higher entropy indicates greater diversity of amino acid usage in LCRs, whereas lower entropy indicates areas characterised by a small subset of residues. Interestingly, AlcoR (Mode 1 and Mode 2) and XSTREAM had high median entropy values ranging from 3.0 to 3.8. This indicates that these algorithms are quite permissive, frequently detecting LCRs with significant amino acid variation even though they are labelled as low-complexity, thus covering a larger range of sequence types. Dotplot and SEG strict, however, showed much lower median entropy values, around 1.5 to 2.0, reflecting strong selection bias towards highly compositionally simple regions composed of a limited number of amino acids. SEG default and intermediate modes, and LCRFinder, exhibited intermediate entropy profiles, finding a combination of strongly biased and moderately heterogeneous LCRs. The fLPS suite of tools (fLPS, fLPS strict, fLPS 2.0, and fLPS 2.0 strict) disclosed wide entropy distributions with comparatively high medians (3.0–3.5), indicative of their design sensitivity to both narrowly and widely biased low-complexity sequences. T-REKS had moderate entropy values and strong variation, reflecting flexible detection by sequence periodicity. Overall, these results establish that SEG, strict and Dotplot are very stringent and prefer plain compositional patterns, while AlcoR, XSTREAM, and the fLPS variants use more general definitions of low complexity, covering a larger set of sequence complexities.

### The detection consensus of the independent methods hints towards regions of homorepeats

We conducted a comprehensive analysis of low-complexity regions (LCRs) by examining how the compositional characteristics of the regions commonly detected across tools varied with the number of independent computational tools that identified them, thereby assessing the relationship between detection consensus and sequence properties (**Fig. 2**). This analysis revealed a striking continuum in both sequence diversity and complexity. LCRs identified by one or more limited tools (in most cases 1–3) displayed extensive heterogeneity of amino acid composition, with often a wide variety of residues and intricate motif types. Such regions had higher Shannon entropy and lower purity values of sequences, which correspond to no strong residue usage and a larger level of randomness or compositional richness. Such features indicate that these LCRs reside outside the canonical definition of low complexity, possibly including context-specific or structure-sensitive sequences. The more tools identified a given LCR, the more significant the shift in compositional features became. The incidence of simple, compositionally skewed motifs—especially homopolymers like polyalanine (A), polyleucine (L), and polyserine (S), and simple dipeptide and tripeptide repeats like AA, SS, and GG—increased markedly. LCRs consistently detected by 10 or more tools had extremely low entropy and highly pure values (median values near 0.8–1.0), indicating a strong enrichment for homorepetitive and compositionally extreme sequences. This pattern was particularly widespread in areas identified by all 13 tools, which predominantly represented typical LCRs with little sequence complexity. The convergent identification of such areas probably owes to a mix of algorithmic bias—given that most tools are tuned to identify strong compositional skew—and the biological importance of such simple, repetitive motifs in protein function, aggregation, or phase separation. Significantly, this gradient highlights the value of methodological heterogeneity in LCR detection: whereas high-consensus regions can be genuine biological hot spots of compositional bias, a strict consensus-based approach will discard compositionally dense or unusual LCRs. Therefore, combining findings from both strict and relaxed detection thresholds from different algorithms allows a more inclusive capture of the LCR landscape, encompassing both highly repetitive and subtly biased sequences with functional significance.

**Figure 2.**
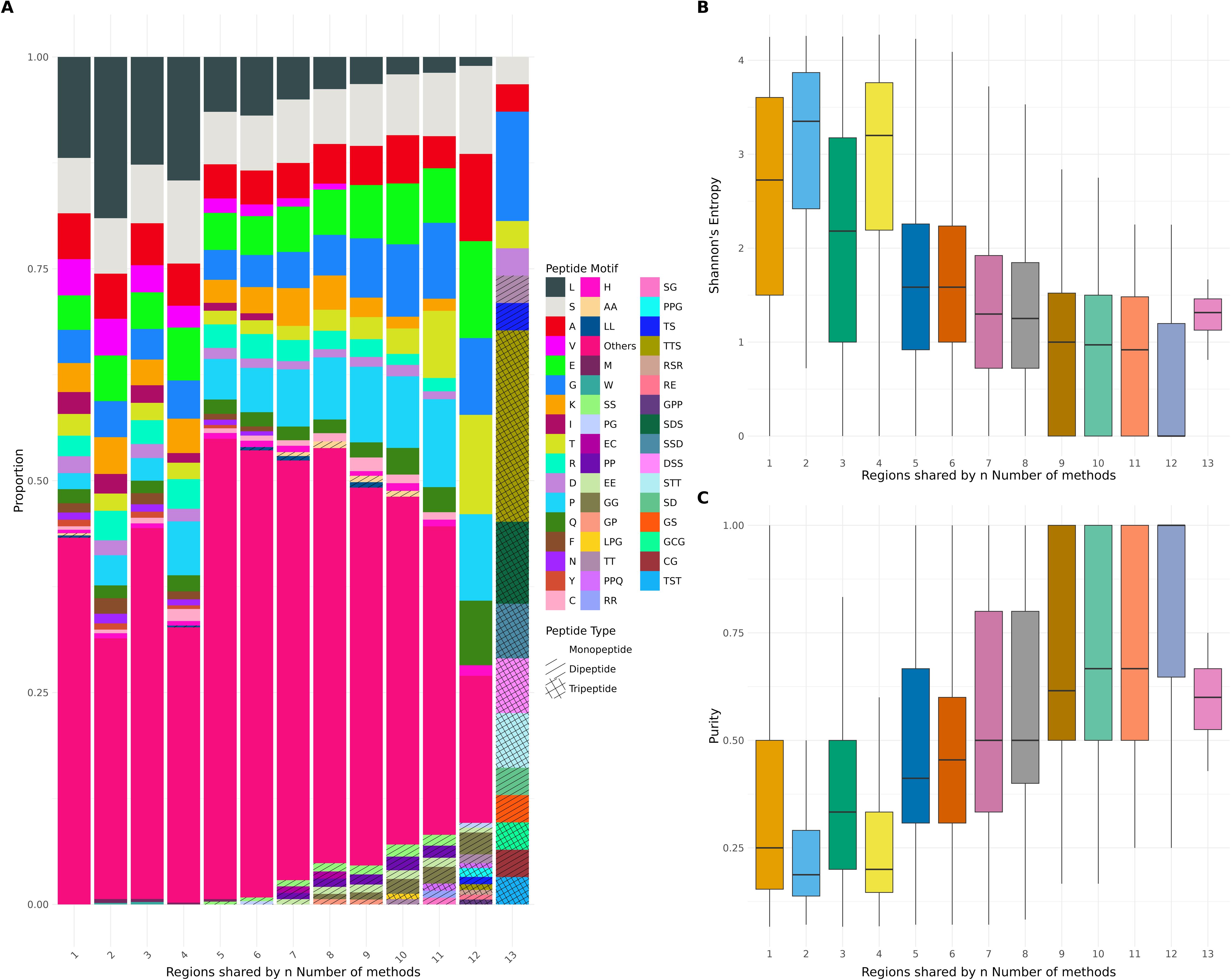
Amino acid composition bias among LCRs detected by different methods. The figure presents the relative frequency of each amino acid within the detected LCRs across all methods. The x-axis lists the 20 standard amino acids, and the y-axis shows the proportion (in percentage) of each residue. Each colour-coded bar corresponds to a specific detection tool, as indicated in the legend. Methods favouring compositionally biased or repetitive regions exhibit overrepresentation of certain residues (e.g., alanine, glycine, serine, or proline), while others show more balanced amino acid distributions. This comparison highlights methodological differences in capturing sequence compositional biases.

### Low-complexity sequence space is progressively constrained by method agreement

In **Fig. 2**, a pattern of decrease in entropy and increase in sequence purity is observed, with an increase in the consensus of the number of methods used. Observing this trend, efforts were made to visualise the LC sequence space by increasing the consensus of the number of methods used (where *k* = number of methods). In **Suppl. Fig. 1**, density contours reveal that all *k* tiers lie along a common diagonal manifold in LC space, but increasing *k* progressively sharpens this distribution. Low-consensus regions occupy diffuse, multimodal densities, whereas high-consensus regions collapse into compact peaks. Thus, consensus does not redefine low complexity, but increasingly restricts it to a narrow, well-defined subset.

In **Suppl. Fig. 2**, centroid trajectories across *k* values demonstrate a coherent directional flow: increasing *k* corresponds to increasing mutation tolerance and decreasing compositional dominance. Variability is high at low *k*, reflecting method-specific definitions, but narrows substantially at high *k*, indicating convergence toward a shared LC regime. This monotonic progression implies that consensus effectively filters out borderline cases, retaining only regions with strong, method-independent low-complexity signatures.

In **Suppl. Fig. 3**, the cumulative consensus map shows that the highest *k* values do not occur at extremes of compositional bias, but instead cluster in intermediate regions combining moderate dominance with elevated mutation rates. Regions with low mutation and low dominance remain largely unoccupied, while extreme repeats show limited consensus. This pattern indicates that methodological agreement is highest for LC regions balancing compositional bias and sequence variability, rather than for strictly repetitive motifs.

In **Suppl. Fig. 4**, stratification by consensus tier (*k*) reveals that agreement among LC detection methods is not uniform across sequence space. Low *k* tiers (1–3) span broad regions with moderate mutation and substantial amino-acid dominance, indicating that many methods independently flag compositionally biased segments. As *k* increases, the occupied space contracts and shifts toward lower mutation percentages and higher dominance. High-consensus regions (*k* ≥ 9) are rare and confined to highly heterogeneous sequences, suggesting that only LC regions are robustly recognised across algorithms.

### The retention rate of LCRs predicted by independent methods drops with increasing sequence purity

The purity-retention profiles in **Fig. 3** indicate unique behaviours for each LCR detection tool with increasing compositional bias (purity). LCRFinder is one of the most uniform tools at all purity levels, retaining high values even at very high purity limits. Its performance curve is one of gradual decline, indicating strong sensitivity to highly repetitive and homopolymeric areas. Dotplot also retains relatively well at moderate to high purity levels but starts to fall more precipitously after ∼75% purity, suggesting good but slightly less sensitivity than LCRFinder. SEG strict exhibits excellent retention across the entire purity range, even better than most tools at high purity levels, and is therefore especially well suited to identifying highly compositionally biased regions. Its intermediate and default forms (SEG intermediate and SEG) have similar patterns but with drop-offs in retention occurring earlier, indicating a compromise between sensitivity and specificity. T-REKS has a moderate percentage of sequences retained at low-to-mid purity levels but begins a consistent decline after 50% purity, indicating that it prefers moderately biased areas over very pure homorepeats. The fLPS family of programs displays a definite hierarchy according to stringency of parameters. fLPS and fLPS 2.0 have similar retention profiles, trapping a wide range of LCRs with moderate purity, but both fall off steeply at purities >50–60%. Their more stringent versions (fLPS strict and fLPS 2.0 strict) have significantly lower retention at lower purities and fall off steeply, suggesting their stringent bias towards highly repetitive, homopolymeric sequences. This indicates how the same algorithm structure can be adjusted to move the detection spectrum towards either broader or narrower compositional definitions. AlcoR (Mode 1 and Mode 2), on the other hand, maintains a high percentage of LCRs only at low purity levels, with a sharp fall beyond 25–30% purity, and practically no sequences preserved beyond 50% purity. This trend indicates that AlcoR is best suited for identifying complex, compositionally heterogeneous LCRs and is less effective at finding simple homorepeats. Likewise, XSTREAM has one of the most precipitous retention declines of any tool, detecting a wide range of low-purity LCRs but losing the majority of detections with increasing purity. Its curve highlights an extremely strong bias toward identifying complex, heterogeneous regions. Overall, every tool has a distinct reaction to compositional purity increase. Applications such as LCRFinder, SEG strict, and Dotplot are very efficient in identifying plain, repetitive LCRs. Some others, such as AlcoR and XSTREAM, perform well to pick up complex, low-purity areas. XSTREAM was also run with different -m parameters (1 to 5), on the human proteome. However, the results come out to be almost identical for -m values of 2 to 5. Very similar sharp drops in entity retention are observed when the sequence purity is gradually increased (**Suppl. Fig. 9**). Only at -m = 1, almost 38% of the entities are retained, even at 100% sequence purity. [At default mode, the -m parameter is set at 5.] The fLPS and SEG groups provide sensitivity tunability so that flexible deployment may be based on target LCR properties. These biases in individual tools highlight the utility of integrative methods, making use of complementary strengths to derive a more comprehensive LCR picture.

**Figure 3.**
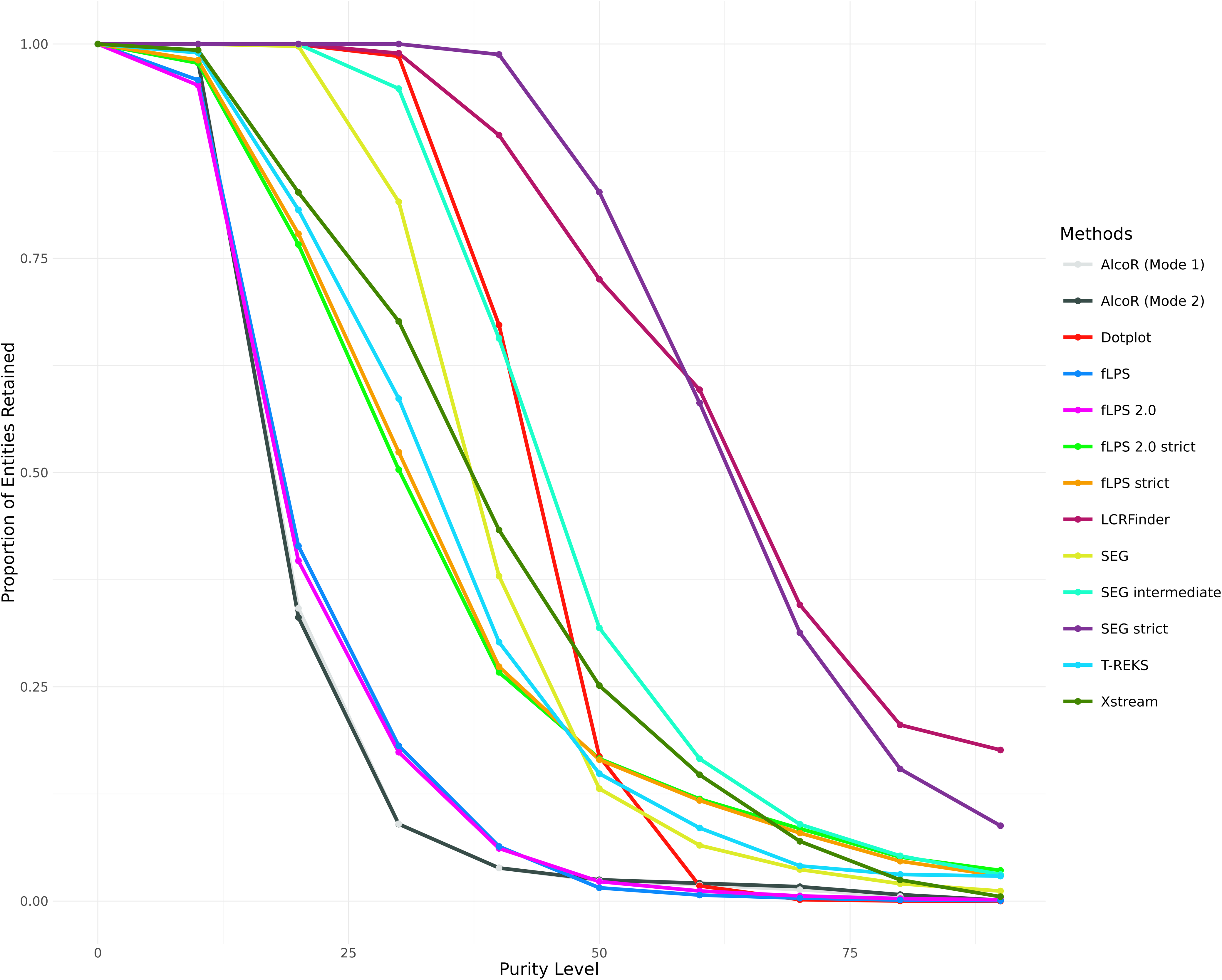
Purity distribution of LCRs detected by different computational methods. This figure shows the variation in LCR purity across detection methods. The x-axis represents the purity level, defined as the percentage of the most dominant amino acid within each LCR, while the y-axis denotes the proportion of total LCRs retained at or above each purity threshold. Each line corresponds to a different method, as indicated by the colour-coded legend. Steeper declines indicate methods that detect fewer highly pure (compositionally biased) regions, whereas gradual slopes reflect identification of more homogeneous LCRs. This comparison emphasises how algorithms vary in their sensitivity to amino acid repetitiveness.

### The similarity between the methods is dependent on their working algorithms

**Fig. 4** illustrates several top LCR detection algorithms which were compared in terms of their pairwise residue-level overlap using the scores in a Jaccard Matrix, providing insight into both their algorithmic foundation and the types of regions they tend to detect preferentially. The conventional SEG method, which identifies regions of low sequence complexity through computation of Shannon entropy in moving windows, shared a moderate degree of overlap (represented by the Jaccard scores) with compositionally skewed tools: 0.1235 with fLPS 2.0 strict, 0.0981 with fLPS 2.0, and 0.0509 with AlcoR. However, its overlap with repeat-detecting tools like T-REKS (0.1399) and Dotplot 0.3706) was quite high, suggesting that SEG could also identify repetitive regions if they possess less complex compositions. fLPS and its updated version, fLPS 2.0 (strict modes enabled), showed very high reciprocal overlap (0.9462 between fLPS 2.0 and fLPS, and 0.6728 between fLPS2 and fLPS 2.0 strict), reflecting their amino acid compositional bias sensitivity. The strict modes, designed to suppress false positives, as expected, overlapped less with the rest of the tools (e.g., fLPS and SEG: 0.099, vs. fLPS 2.0 strict and SEG: 0.1235), reflecting that parameter optimisation influences region identification. AlcoR and AlcoR 2 (AlcoR operated in 2nd mode), from statistical profiles of amino acid usage, presented high inter-method concordance (0.7236), thus validating method reproducibility. Crossmatches with other methods, such as fLPS 2.0 (0.1031) and T-REKS (0.1179), further suggest that AlcoR methods detect both repetitive and compositional components to a limited extent. The Dotplot technique, which identifies self-similarity between sequences through comparisons of overlap windows, showed minimal overlap with most composition-based tools: 0.0346 with AlcoR, 0.0478 with fLPS, and 0.0621 with fLPS 2.0 strict. However, it did have significant overlap with SEG (0.3706) and LCRFinder (0.2557), demonstrating its capacity to identify visually separable repeat-like structures that are also compositionally simple. LCRFinder, detecting highly frequent motifs (with mismatches), was closest in its top overlap to Dotplot (0.2557) and SEG (0.1853), but considerably less so with algorithms like fLPS 2.0 strict (0.0375) or AlcoR (0.0299), indicating its emphasis on detecting local repetitive motifs more than compositionally skewed patterns. T-REKS, as an optimised tool for tandem repeat detection by self-alignment and periodicity verification, exhibited maximum overlap with XSTREAM (0.1469) (also a program for finding repeats), followed by moderate overlap with SEG (0.1399) and Dotplot (0.1241). Overlap of the latter with fLPS (0.0424) and AlcoR (0.1179), however, was relatively low, validating its structural rather than compositional repetition target. XSTREAM, employing suffix trees and pattern matching to identify imperfect, complex repeats, overlapped very little with composition-biased methods such as SEG strict (0.0054) and fLPS 2.0 strict (0.0184), but was most similar to T-REKS (0.1469), which indicates its repeat-oriented nature. Finally, in the stricter forms of SEG (e.g., SEG strict), overlaps were systematically reduced, as found between SEG strict vs. SEG (0.0559) and SEG strict vs. fLPS2 (0.0082), which affirms that strict complexity constraints merely permit the discovery of the most well-defined LCRs. These results evidence clear partitioning of algorithms based on detection principles. Composition-biased tools (fLPS, fLPS2, Alcor, SEG) exhibit high mutual overlap due to their bias for amino acid frequency. Tools like Dotplot, T-REKS, XSTREAM, and LCRFinder, which target repetitive or periodic motifs, exhibit relatively low overlap with composition-biased tools but higher agreement amongst themselves. This divergence illustrates the importance of tool selection according to biological context: if one is looking for detecting statistically enriched amino acid regions, tools like fLPS or Alcor are best; to identify repetitive, periodic or motif-based, T-REKS or XSTREAM are better.

**Figure 4.**
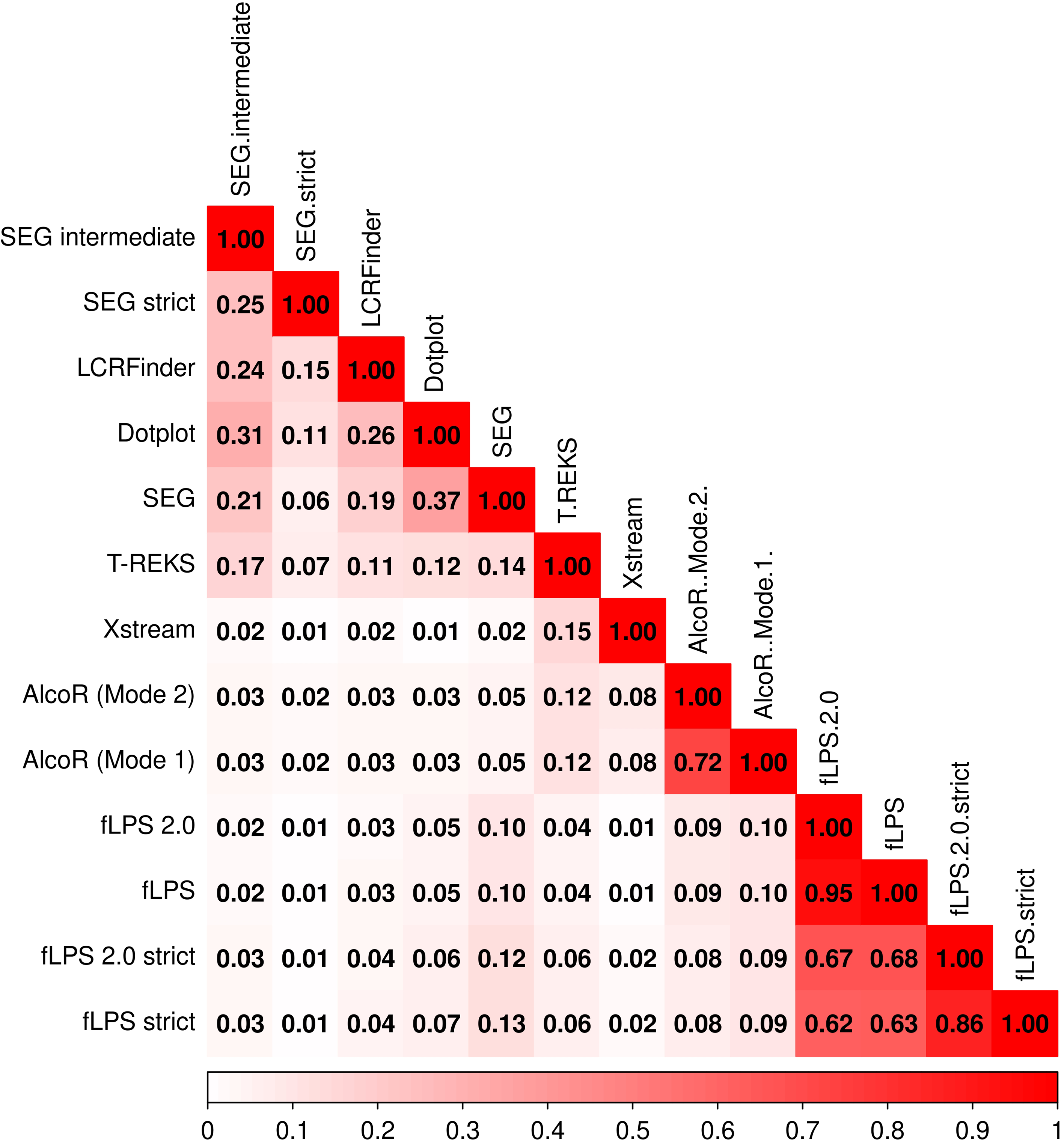
Pairwise overlap and similarity between LCR detection methods. It presents the Jaccard similarity index for each pair of methods, calculated as the intersection divided by the union of detected LCR sets. The colour gradient (white to red) denotes increasing similarity, with diagonal values (1.0) representing self-comparisons. Together, these heatmaps quantify the extent of agreement and divergence among the evaluated detection algorithms.

### True LCR boundaries are fundamentally fuzzy and ambiguous

**Suppl. Fig. 5** and **Suppl. Fig. 6** present probabilistic reconstructions of low-complexity (LC) space across increasing consensus stringency (*k* = 7–12). In both figures, each panel shows a posterior probability surface *P*(LCR∣bin) in mutation–composition space, with black contours indicating ROC-optimised decision boundaries. Across both datasets, increasing *k* produces a systematic contraction of the LC-permissive region toward the low-mutation, high-compositional-bias corner of the space. This trend reflects the fact that higher consensus thresholds preferentially retain sequence segments that are consistently identified as low-complexity by many independent methods, thereby enriching for highly repetitive sequences. Correspondingly, the ROC-optimised probability threshold decreases monotonically with *k*, indicating that increasingly confident LC assignments emerge from progressively smaller regions of sequence space. Importantly, the inferred boundaries are not sharp demarcations, but fuzzy, graded transition zones. The posterior landscapes exhibit smooth probability gradients rather than step-like separations, underscoring that low complexity is not a binary sequence property. Instead, LC behaviour arises from a continuum of sequence architectures governed by compositional entropy and mutational tolerance. Regions near the diagonal boundary—where mutation rate and amino-acid dominance trade off—retain intermediate posterior probabilities and shift gradually with **k**, reflecting ambiguity rather than misclassification. Comparison between PDB-based (**Suppl. Fig. 5**) and DisProt-based (**Suppl. Fig. 6**) maps further highlights that boundary placement depends on the underlying biological reference. DisProt-derived boundaries extend further into higher-mutation regimes, consistent with the presence of experimentally validated disorder that tolerates greater sequence variability, whereas UniProt-derived boundaries are more restrictive.

The GAM (Generative Additive Model) posterior maps derived from DisProt- and PDB-anchored models (**Suppl. Fig. 7**; **Suppl. Fig. 8**) provide a unified view of how low-complexity (LC) space is structured by sequence entropy and compositional bias. Across all panels (A–F), LC propensity is organised along a curved boundary in the two-dimensional space defined by mutation percentage and dominance of the most frequent amino acid. This boundary is not arbitrary: it reflects a tradeoff whereby increasing mutational heterogeneity necessitates reduced compositional bias, and vice versa. Regions combining high mutation with high dominance are systematically excluded, indicating sequence architectures that are rarely compatible with either curated regions missing in PDB structures or experimentally validated disorder (DisProt). A key trend emerges across panels as the minimum method consensus constraint (k_min) increases. At lower k values (A–C), the LC space is expansive and irregular, encompassing short, compositionally mixed regions that satisfy permissive disorder criteria. As k increases (D–F), the LC space contracts progressively, and the decision boundary becomes smoother and more monotonic. This shrinkage indicates that LC regions detected with consensus across more methods must satisfy stricter entropic constraints, filtering out marginal or noisy configurations and retaining only sequence architectures with robust disorder signatures. Notably, DisProt-derived boundaries are consistently more restrictive than PDB-derived ones, underscoring the tighter constraints imposed by experimental validation relative to missing residue-based curation.

Together, these figures demonstrate that low-complexity (LC) boundaries are not sharp decision surfaces but probabilistic envelopes embedded in sequence-information space, whose shape depends on both reference background sequences and scale. Apparent LC space emerges gradually as compositional bias strengthens and effective sequence entropy decreases, rather than at a single, universal cutoff. At the same time, LC space is intrinsically scale-dependent: increasing the minimum method consensus constraint (k) systematically contracts this space and smooths its geometry, reflecting progressively stricter entropic requirements for regions detected across a greater number of methods. The convergence toward stable, smooth boundaries at higher k does not resolve LC into a uniquely defined class, but instead highlights that “true” LCR boundaries remain fundamentally fuzzy and ambiguous. Collectively, these results support an information-theoretic view of intrinsic disorder in which allowable LC architectures occupy a constrained, annotation-dependent envelope governed by the balance between entropy and compositional bias, rather than a fixed or discrete notion of low complexity.

### LCR complexity profiles of the methods show varying degrees of clustering

To further evaluate the behaviour of various low-complexity region (LCR) detection tools, we analysed the two-dimensional distribution of percentage mutation and percentage most common amino acid inside predicted LCRs, employing the Low Complexity triangle model depicted in **Fig. 5**. This triangle divides sequence space into areas corresponding to homorepeats (top), long imperfect repeats (bottom left), low-complexity regions (centre), and high-complexity sequences (bottom right). The heatmaps of density for each of the tools demonstrate their respective detection biases and sensitivities in this compositional and mutational landscape.

**Figure 5.**
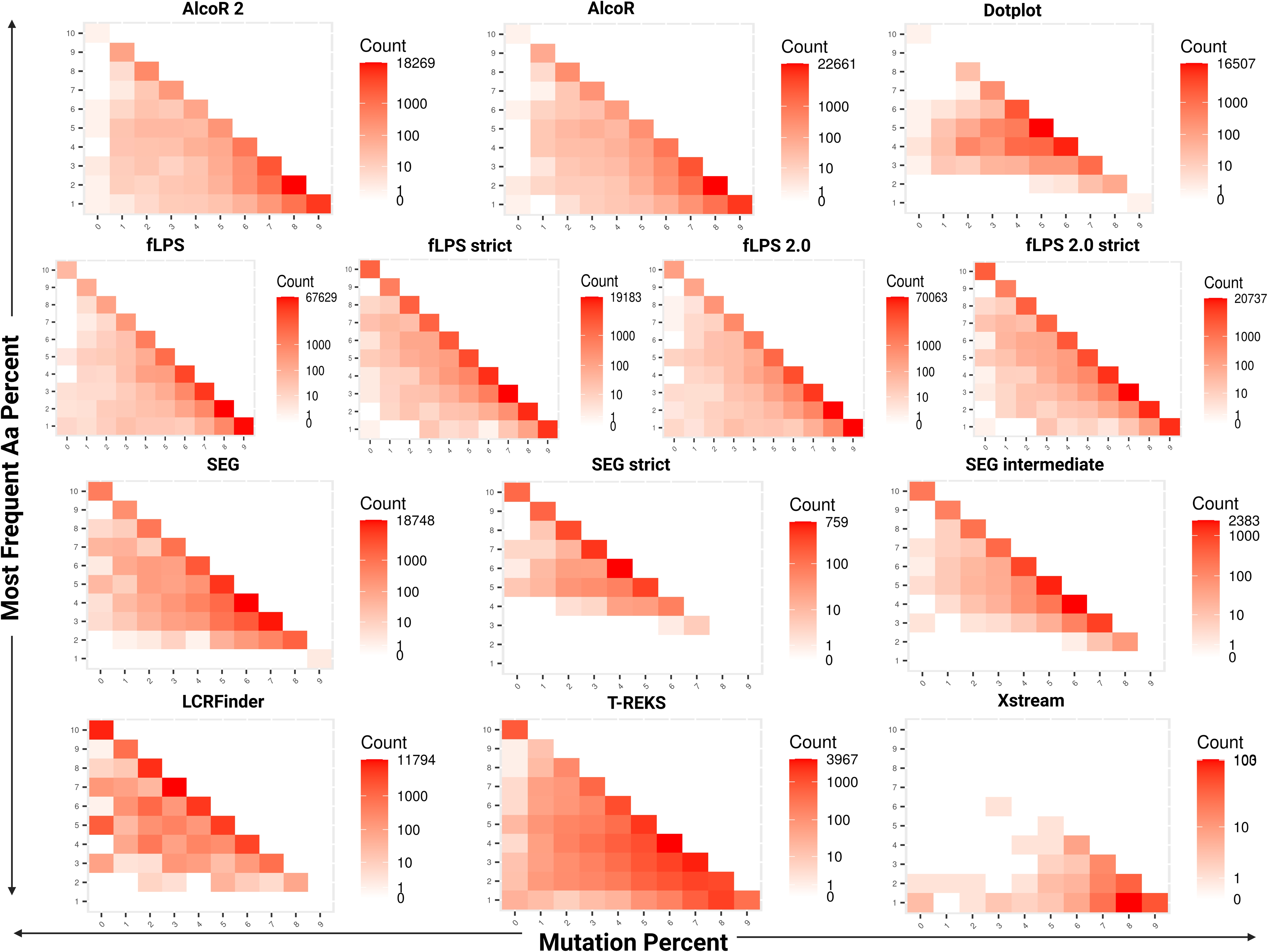
Relationship between amino acid composition and mutation percentage in LCRs across detection methods. Each panel corresponds to one detection method and illustrates the joint distribution of amino acid composition bias and sequence mutation percentage. The x-axis represents mutation percentage (fraction of variable residues within the LCR), and the y-axis shows the percentage of the most frequent amino acid. The colour gradient (white to red) indicates the number of LCRs within each bin, as shown by the colour key. The diagonal trend visible across most methods indicates that higher amino acid dominance is typically associated with lower mutation percentages, reflecting stronger compositional homogeneity in repetitive regions.

AlcoR and AlcoR2 both showed high concentrations of dark red signal in the bottom-right corner of the triangle, which represents sequences with high rates of mutation and comparatively low amino acid bias. These trends show that both tools often predict relatively noisy and complex areas as LCRs. Their intrinsic algorithms, which take compositional skew and k-mer frequency metrics into account, seem to be geared towards detecting "fuzzy" LCRs—regions that are far from perfect repeats but exhibit faint compositional anomalies. Although this renders AlcoR and AlcoR2 very sensitive, it also implies a liberal detection mechanism that might sacrifice specificity for the sake of coverage, detecting areas that other methods would filter out as not having enough compositional or structural simplicity.

Dotplot, on the other hand, showed a steeper diagonal pattern in the centre of the triangle. This represents an emphasis on moderately biased, moderately mutated sequence regions—typically regions with classical LCRs. The algorithm’s use of visual self-similarity heuristics causes it to be more discriminating in its detection profile than AlcoR and to have very little detection at high-complexity sequence space. It can avoid overprediction without sacrificing the detection of many low-complexity patterns that are varied in nature.

The fLPS family, comprising fLPS, fLPS strict, fLPS 2.0, and fLPS 2.0 strict, showed universal detection, with darker areas stretching from the top (high amino acid bias, i.e., homorepeats) towards the centre. They are very good at detecting both highly biased homopolymeric tracts and moderately imperfect LCRs, reflecting their compositional filtering and scoring flexibility. But the fact that the signal is seen in regions of intermediate mutation implies that the default (non-strict) variants of fLPS are, to some extent, permissive. Their strict counterparts, by contrast, limit predictions to more compositionally extreme and less mutated parts, raising specificity at the risk of excluding important biological variation.

SEG and its variants offered a wide range of detection. The default SEG tool exhibited dark signals from the lower-left to the triangle’s centre, indicating good ability to detect long imperfect repeats and biased low-complexity sequences. SEG’s entropy-based scoring accommodates this flexibility. The intermediate and strict versions progressively lowered coverage, with SEG strict narrowly examining areas of high compositional bias and low mutation—largely homorepeats—exhibiting greater stringency at the cost of breadth.

LCRFinder exhibited moderate dark red signals along the left-hand side of the triangle, indicating a bias towards sequences with moderate amino acid bias and relatively low mutation. This trend indicates that LCRFinder balances compositional bias and structural complexity. Its detection approach, which incorporates Shannon entropy with expected-vs-observed residue frequencies, enables it to spot a variety of biased and partially repetitive areas without generating false positives in highly complex or disordered regions—showing improved specificity over AlcoR.

T-REKS, employing a k-means clustering strategy optimised for tandem repeat detection, was found to be located primarily in the central part of the triangle, with a slight bias toward sequences with moderate bias and low-to-moderate mutation. It steers clear of highly complex and highly biased areas, as designed to identify structured, periodic LCRs. T-REKS is slightly more permissive of mutation than LCRFinder, though it still efficiently eliminates high-complexity noise.

XSTREAM showed the most selective profile, with scattered but localised dark red signals restricted to areas of low mutation and moderate amino acid bias. This suggests a strong preference for clean, well-ordered tandem arrays with few indels or compositional anomalies. Its high stringency and low sensitivity make it a good choice for identifying only the most canonical and structurally uniform repeats.

Overall, these complexity heatmaps highlight that LCR detection is strongly tool-dependent. Compositional tools like fLPS, AlcoR, and SEG are optimally suited for the detection of amino acid enrichment or disorder-driven regions, whereas structure-aware tools like T-REKS and XSTREAM emphasise periodicity and sequence regularity. Hybrid methods like LCRFinder provide greater flexibility by combining compositional and statistical aspects. Further, in all the tools, strict or conservative modes always sacrifice sensitivity for accuracy, possibly eliminating noisy but biologically relevant LCRs. These findings underscore the need to choose a suitable detection strategy depending on the particular biological question or sequence context: there is no one tool that can catch all of the diversity of low-complexity sequence space.

### The methods exhibit varying degrees of sensitivity and specificity to LCR detection

**Fig. 6** illustrates performance trends (TPR and FPR) of different LCR detection tools over four variables: gene length (A), number of LCRs per gene (B), LCR coverage percentage (C), and LCR: Gene Entropy Ratio (D). Many common trends of increase and decrease can be noted. Across all the panels, fLPS and fLPS 2.0 (strict variants) exhibit a consistently higher TPR as the gene length, number of LCRs, coverage, and entropy ratio increase. This means that their sensitivity increases with a stronger or more abundant LCR signal. Their FPRs also significantly increase, particularly in Panels A and D, indicative of more false positives with growing complexity, especially for long genes or genes with high entropy. Dotplot and SEG (intermediate and default) have constantly high TPRs on all variables, save for slight fluctuations. Their FPRs are relatively low and constant, but increase with increased length and higher entropy (Panels A and D). They all prove stable over various conditions, with Dotplot producing an overall stability of performance that is optimal. SEG strict is unique in demonstrating a relatively flat TPR trend, being lower than its more relaxed variants but with an exceptionally low and stable FPR under all conditions. The trend supports its application as a high-specificity tool, particularly when minimising false positives. LCRFinder and T-REKS exhibit modest increases in TPR with growing LCR features, while keeping FPR low or moderately increasing, indicating balanced but less stringent detection. By contrast, AlcoR (both modes) and XSTREAM exhibit flat or slightly decreasing TPRs under most conditions, reflecting poor sensitivity despite increasingly prominent LCR features. Their FPRs are low, indicating conservative detection that presumably results in false negatives rather than true specificity. In general, most tools exhibit a positive correlation between TPR and rising LCR-related features, but few (specifically, Dotplot and SEG variants) are able to restrain the increase in FPR, and hence are more suitable for precise LCR detection in a wide range of genomic contexts.

**Figure 6.**
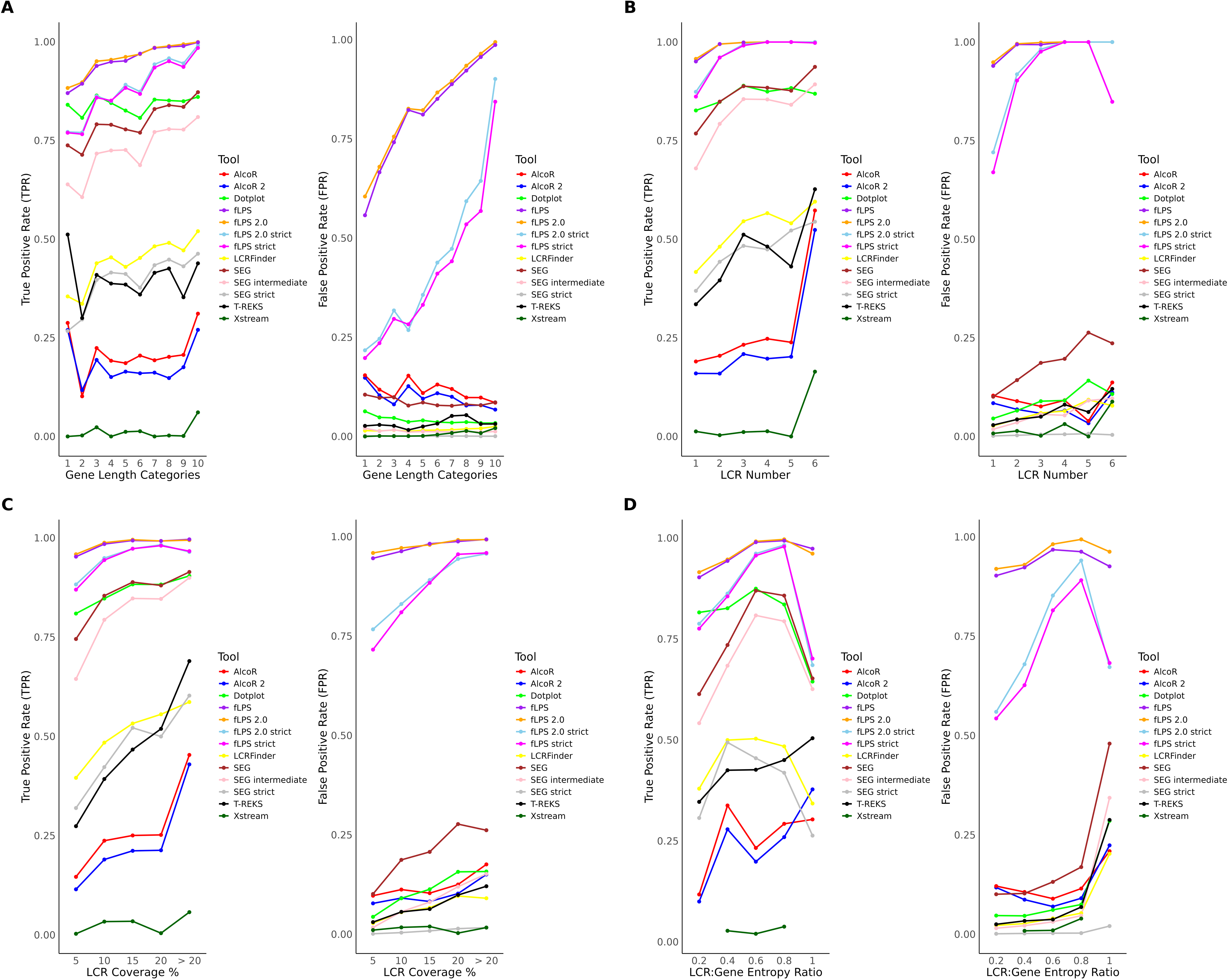
Evaluation of LCR detection performance across sequence and compositional parameters. This figure assesses the performance of different tools in detecting simulated LCRs under varying conditions. True positive rate (TPR; left subpanels) and false positive rate (FPR; right subpanels) are plotted against four parameters: **(A)** gene length categories, **(B)** number of LCRs per gene, **(C)** LCR coverage percentage, and **(D)** the LCR-to-gene entropy ratio. The x-axes represent increasing parameter values, while the y-axes show detection rates. Distinct colours denote individual tools, as indicated in the legend. Higher TPRs indicate more sensitive detection, whereas lower FPRs reflect higher specificity. These comparisons reveal how each method’s performance varies with sequence length, LCR density, coverage, and compositional complexity.

A complete performance comparison of an extensive number of low-complexity region (LCR) detection tools (as shown in **Fig. 7**) —based on an average true positive rate (TPR) and false positive rate (FPR) across a broad set of conditions—reveals that different sensitivity-specificity trade-offs exist between methods. Comparisons were made using a variety of parameters, including gene length, LCR per gene, coverage percentage of the LCRs, and LCR: Gene Entropy Ratio, and hence constituted a rigorous test of the consistency of each methodology under varying genomic circumstances. Among all the tools, fLPS and fLPS 2.0 (both of their strict modes) consistently registered the highest TPRs and exhibited their natural strength for detecting LCRs even at tough and heterogeneous cases. Due to this, they are incredibly efficient in obtaining real LCRs under any typical conditions. But this sensitivity comes with a cost: they also possessed some of the highest FPRs, particularly when gene length, LCR density, and entropy ratio were increasing. This means that while these programs are sensitive to the detection of diverse LCRs, they would overpredict, reducing their specificity by falsely labelling non-LCR regions. SEG, especially default and intermediate modes, and Dotplot, respectively, showed the best possible balance of sensitivity and specificity. Both tools produced high TPRs with comparatively low FPRs, reflecting efficient actual LCR detection with very few false positives. Surprisingly, Dotplot ranked as the best-performing tool overall, consistently outperforming all other methods by rendering correct detection with the least number of false positives. SEG strict, while possessing lower TPRs, had one of the lowest FPRs across all conditions and, therefore, is an excellent choice for high-specificity use where false positives must remain low. LCRFinder and T-REKS were moderately sensitive and had low false positives, indicating a conservative but quite reasonable performance. They both do well in obtaining a good trade-off for overall LCR detection, especially when there is a desire for precision along with some inclusivity. Conversely, AlcoR (default and alternative mode) and XSTREAM all reported low TPRs with low FPRs, likely due to their extremely conservative detection thresholds. This is in agreement with a tendency to under-detect true LCRs and hence be suboptimal for complete LCR identification, particularly in high LCR diversity or compositional complexity datasets. Together, this comparison indicates that there is no single tool that is better all around. These technologies, like fLPS and fLPS 2.0, are appropriate for thorough and sensitive searches but are prone to generating false positives. SEG and Dotplot are more balanced and stable in their performance and are top contenders as efficient, general-purpose LCR detectors. If high specificity is required, SEG strict is the most precise tool. Ultimately, the choice should be made on the basis of biological context and tolerance for false positives, as performance can be extremely variable as a function of complexity and type of sequence data.

**Figure 7.**
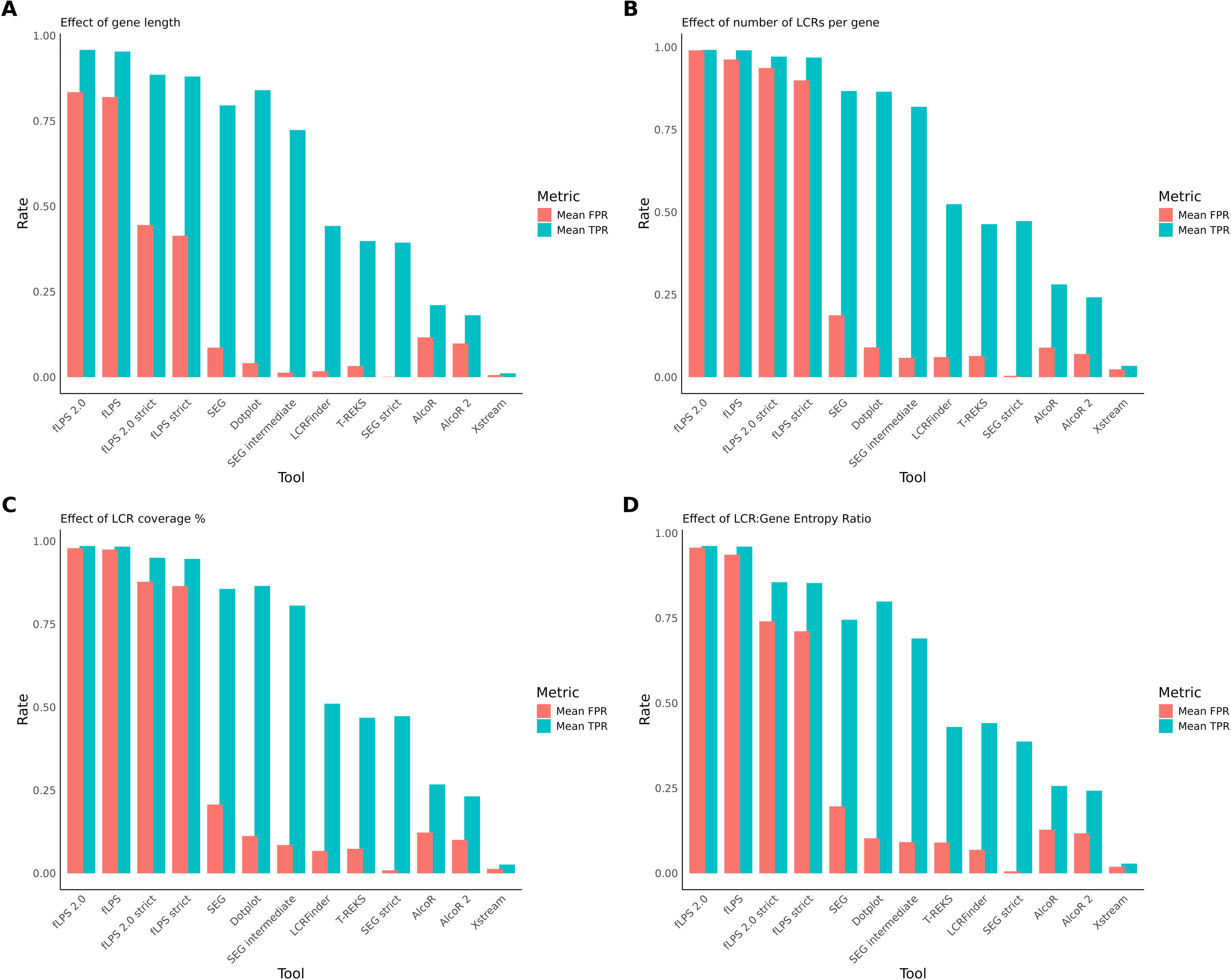
Comparative summary of average detection performance metrics across tools. This figure presents the mean true positive rate (TPR, cyan) and mean false positive rate (FPR, pink) for all detection methods across the same four parameter categories shown in Figure 7. **(A)** shows the effect of gene length, **(B)** depicts the effect of the number of LCRs per gene, **(C)** shows the effect of LCR coverage percentage, and **(D)** represents the effect of the LCR-to-gene entropy ratio. The x-axis lists all detection tools, and the y-axis indicates normalised rate values on a 0–1 scale. This integrated summary provides a direct comparison of each method’s overall sensitivity and specificity across multiple structural and compositional parameters.

**Figure 8.**
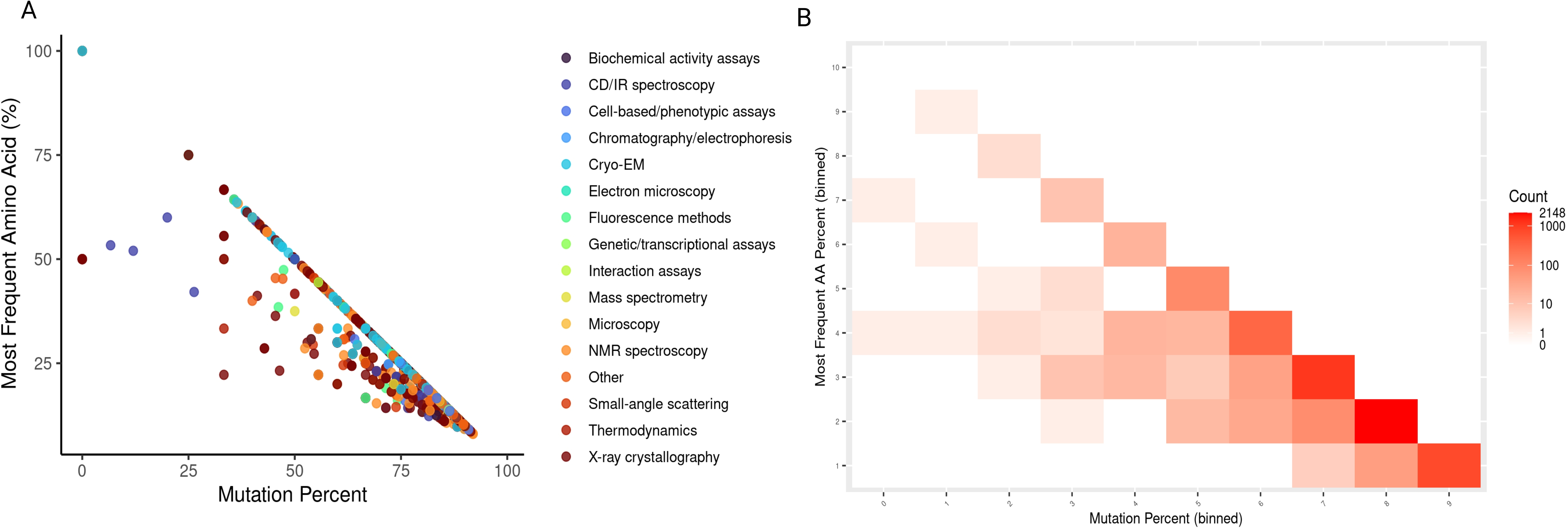
Visualising the complexity of structurally disordered regions. **(A)** Scatter plot of DisProt-annotated intrinsically disordered regions showing mutation percentage versus the most frequent amino acid percentage. Points are colored by experimental method. Regions from diverse techniques occupy a common triangular envelope, indicating a conserved sequence-complexity constraint independent of annotation method. **(B)** Heatmap showing the distribution of PDB regions lacking resolved coordinates projected onto mutation percentage (x-axis) and most frequent amino acid percentage (y-axis). Colour intensity indicates segment counts (log-scaled). Missing residues are enriched at high mutation percentages and low compositional dominance, consistent with highly heterogeneous sequence regions.

## DISCUSSION

Our systematic benchmarking of low-complexity region (LCR) detection tools across the human proteome highlights several important insights with broad implications for protein sequence annotation, functional inference and evolutionary interpretation. First, the clear divergence in length-, coverage- and entropy-based detection profiles between methods underscores that LCRs are not a monolithic class of sequence features, but rather a heterogeneous collection of compartments that differ in size, compositional bias, repetitiveness and biological context. For example, our finding that tool A (composition-bias tuned) and tool B (repeat-periodicity tuned) capture largely non-overlapping LCR sets echoes that “statistical measures alone cannot capture all structural aspects of LCRs”.^26^ Second, our observation that LCRs common to multiple tools tend to be longer, more repetitive (i.e., lower Shannon entropy) and more consistently detected suggests that detection consensus may by itself be a useful proxy for functional relevance. In that regard, previous studies have shown that LCRs located at protein termini tend to be associated with increased numbers of binding partners.^54^ Our multi-tool consensus may preferentially capture, enrich LCRs with stronger structural or interaction-driven constraints, although direct functional validation will be required to establish this link. Third, the correlation heatmaps and Jaccard similarity analysis revealed tight clustering of methods that use similar algorithmic lenses (e.g., composition vs. repeat detection), which aligns with previous evolutionary analyses. For instance, it has been shown that LCR conservation depends on functional context and subcellular localisation, suggesting that different algorithms may preferentially detect distinct sequence subclasses. ^55^ Fourth, our entropy and purity analyses contribute to the emerging view of LCRs as functional modules rather than being uniformly neutral or non-functional. It has been found that many LCRs within disordered regions may adopt secondary structures (e.g., helices). ^56^ Our results show that lower-entropy (more repetitive) LCRs are preferentially retained across multiple detection methods. This enrichment may reflect a combination of biological constraint and intrinsic sequence properties, but does not imply that low-entropy regions are universally maintained by evolutionary selection. Indeed, such regions can arise through stochastic processes such as replication slippage, and only a subset is likely to acquire or retain functional relevance^11^. Fifth, the observed association of LCR-containing proteins with lower protein abundance but higher transcript abundance ^57^ offers a useful context for interpreting our coverage-based findings, suggesting that LCRs may influence translational efficiency or post-translational stability rather than transcript production per se. Our work also highlights specific methodological recommendations. A unified reporting framework that includes length-bin distribution, coverage percentage, amino-acid composition, Shannon entropy and consensus frequency (number of tools detecting) offers richer annotation of LCR-calls than simple “present/absent”.^58^ In practical terms, our results suggest that functional studies of LCRs (e.g., phase separation, promiscuous interactions, aggregation) should prioritise LCRs that are long (> 50 aa), have high-coverage within a protein (> 20%), and are detected by multiple independent algorithms – because these appear enriched for lower entropy and higher compositional purity. This prioritisation strategy is consistent with previously observed links between repetitive or compositionally biased sequences and structural or functional properties, while remaining exploratory in nature^59^. From an evolutionary perspective, our detection-consensus clustering provides a quantitative basis for assessing LCR “families” of detection methods, which may reflect underlying evolutionary modes of LCR origin (e.g., slippage, duplication, expansion). In particular, poor correlation between protein and DNA low-complexity sequences ^60^ suggests that many protein LCRs reflect post-translational or structural constraints that are invisible at the DNA level. Platforms such as PlaToLoCo provide a useful aggregation framework for contextual comparison of low-complexity region (LCR) annotations across multiple detection methods. However, because PlaToLoCo largely integrates outputs from the same underlying algorithms evaluated here, its annotations are not independent and should not be treated as ground truth. Accordingly, such platforms are best viewed as exploratory or comparative resources rather than benchmarks for method validation.^28^ Our analyses further suggest that both structurally unresolved regions in the Protein Data Bank and experimentally validated intrinsically disordered regions are governed by shared sequence-level constraints linking amino-acid composition, mutational tolerance, and conformational flexibility. Regions with high sequence heterogeneity are more likely to lack resolved structural coordinates, consistent with the long-standing view that extensive compositional diversity disfavors stable structure formation. ^49,61^At the same time, the presence of missing residues within compositionally biased segments indicates that low complexity alone is insufficient to explain structural absence, implicating additional factors such as dynamics and experimental context.^62,63^ Experimentally curated disorder occupies a constrained, method-independent region of sequence-complexity space, supporting the idea that intrinsic disorder reflects regulated and evolutionarily maintained sequence architectures rather than random compositional noise.^30,64,65^ In conclusion, our benchmarking and visualisation framework advances the annotation of low-complexity regions by moving beyond single-tool outputs to a multi-metric consensus model. It provides a roadmap for prioritising biologically meaningful LCRs and encourages more nuanced interpretation in studies of sequence complexity, structure and function.

## LIMITATIONS AND FUTURE DIRECTIONS

While the present benchmarking framework provides a comprehensive comparative analysis of existing LCR detection tools, several limitations warrant consideration. First, the study focuses exclusively on the human proteome, and LCR features may vary substantially across phylogenetic lineages, particularly in organisms with extreme GC content or compact genomes. Expanding this framework to include diverse taxa could elucidate evolutionary trends in LCR expansion and constraint. Although we benchmarked 8 major tools, emerging methods (e.g., machine-learning based) might detect additional classes of LCRs, and our human-proteome focus may limit generalisation to other lineages (e.g., prokaryotes, pathogens) where LCR dynamics differ ^66^. Second, our evaluation is based on algorithmic and compositional metrics rather than experimentally validated functional data. Integrating proteomic, structural, or biophysical evidence—such as phase-separation propensity or disorder–order transitions—would strengthen the biological interpretation of computationally predicted LCRs. Third, we note that restricting the dataset to a single representative isoform may underrepresent LCRs that are preferentially introduced or expanded through alternative splicing, particularly in intrinsically disordered terminal or linker regions. However, this choice was made to enable consistent cross-method benchmarking and to minimise redundancy-driven biases. Fourth, missing PDB residues reflect both intrinsic disorder and experimental limitations, and therefore cannot be interpreted as a direct proxy for disorder. Along with that, the analysis remains sequence-centric and does not explicitly account for cellular context, post-translational modifications, or interaction-induced folding, which may modulate disorder in vivo. This calls for sophisticated and well-planned experimental biology approaches. Finally, emerging machine-learning and deep-learning models trained on multi-parameter datasets could capture complex relationships between sequence composition, disorder, and functionality that remain inaccessible to traditional heuristic tools. Future developments should therefore aim toward a hybrid consensus paradigm, combining statistical, compositional, and predictive approaches to achieve more precise, biologically meaningful LCR annotation.

## ETHICS

This work did not require ethical approval from a human subject or animal welfare committee.

## DATA ACCESSIBILITY

The data and relevant code are available in GitHub (https://github.com/AnirjitChatterjee/Benchmarking_LCR_Tools.git).

## AUTHORS’ CONTRIBUTIONS

A.C.: conceptualisation, data curation, formal analysis, investigation, methodology, validation, visualisation, writing— original draft, writing—review and editing; N.V.: conceptualisation, funding acquisition, project administration, resources, supervision, writing —original draft, writing—review and editing.

Both authors gave final approval for publication and agreed to be held accountable for the work performed therein.

## CONFLICT OF INTEREST DECLARATION

We declare we have no competing interests.

## AI USE DECLARATION STATEMENT

AI has been used for paraphrasing the text, as permitted by the Journal Guidelines.

## FUNDING

This article was funded by the Department of Biotechnology, Ministry of Science and Technology, India (grant no. BT/11/IYBA/2018/03) and Science and Engineering Research Board (grant no. ECR/2017/001430) provided funds for procuring computational resources (i.e. Har Gobind Khorana Computational Biology cluster) used.

## Notes

https://github.com/AnirjitChatterjee/Benchmarking_LCR_Tools/tree/main

### Competing Interest Statement

The authors have declared no competing interest.

## REFERENCES

1. Levinson, G. & Gutman, G. A. Slipped-strand mispairing: a major mechanism for DNA sequence evolution. Mol. Biol. Evol. 4, 203–21 (1987).

2. Ellegren, H. Heterogeneous mutation processes in human microsatellite DNA sequences. Nat. Genet. 24, 400–402 (2000).

3. Fondon, J. W. & Garner, H. R. Molecular origins of rapid and continuous morphological evolution. Proceedings of the National Academy of Sciences 101, 18058–18063 (2004).

4. Harbi, D., Kumar, M. & Harrison, P. M. LPS-annotate: complete annotation of compositionally biased regions in the protein knowledgebase. Database (Oxford). 2011, baq031 (2011).

5. Tautz, D., Trick, M. & Dover, G. A. Cryptic simplicity in DNA is a major source of genetic variation. Nature 322, 652–6.

6. Kajava, A. V. Tandem repeats in proteins: from sequence to structure. J. Struct. Biol. 179, 279–88 (2012).

7. Mier, P. et al. Disentangling the complexity of low complexity proteins. Brief. Bioinform. 21, 458–472 (2020).

8. Golding, G. B. Simple sequence is abundant in eukaryotic proteins. Protein Sci. 8, 1358–61 (1999).

9. Mier, P. & Andrade-Navarro, M. A. Assessing the low complexity of protein sequences via the low complexity triangle. PLoS One 15, e0239154 (2020).

10. Lee, B., Jaberi-Lashkari, N. & Calo, E. A unified view of low complexity regions (LCRs) across species. Elife 11, (2022).

11. Radó-Trilla, N. & Albà, Mm. Dissecting the role of low-complexity regions in the evolution of vertebrate proteins. BMC Evol. Biol. 12, 155 (2012).

12. Karlin, S., Brocchieri, L., Bergman, A., Mrazek, J. & Gentles, A. J. Amino acid runs in eukaryotic proteomes and disease associations. Proc. Natl. Acad. Sci. U. S. A. 99, 333–8 (2002).

13. King, D. G., Soller, M. & Kashi, Y. Evolutionary tuning knobs. Endeavour 21, 36–40 (1997).

14. Gemayel, R., Cho, J., Boeynaems, S. & Verstrepen, K. J. Beyond Junk-Variable Tandem Repeats as Facilitators of Rapid Evolution of Regulatory and Coding Sequences. Genes (Basel). 3, 461–480 (2012).

15. Newton, A. H. & Pask, A. J. Evolution and expansion of the RUNX2 QA repeat corresponds with the emergence of vertebrate complexity. Commun. Biol. 3, 771 (2020).

16. Reiner, A., Dragatsis, I. & Dietrich, P. Genetics and Neuropathology of Huntington’s Disease. in 325–372 (2011). doi:10.1016/B978-0-12-381328-2.00014-6.

17. Kedzierski, Ł., Montgomery, J., Curtis, J. & Handman, E. Leucine-rich repeats in host-pathogen interactions. Arch. Immunol. Ther. Exp. (Warsz). 52, 104–12 (2004).

18. Hancock, J. M. & Simon, M. Simple sequence repeats in proteins and their significance for network evolution. Gene 345, 113–8 (2005).

19. Kashi, Y. & King, D. G. Simple sequence repeats as advantageous mutators in evolution. Trends Genet. 22, 253–9 (2006).

20. Lynch, V. J. & Wagner, G. P. Resurrecting the role of transcription factor change in developmental evolution. Evolution 62, 2131–54 (2008).

21. Franzmann, T. M. & Alberti, S. Prion-like low-complexity sequences: Key regulators of protein solubility and phase behavior. Journal of Biological Chemistry 294, 7128–7136 (2019).

22. Chong, P. A., Vernon, R. M. & Forman-Kay, J. D. RGG/RG Motif Regions in RNA Binding and Phase Separation. J. Mol. Biol. 430, 4650–4665 (2018).

23. Kato, M., Lin, Y. & McKnight, S. L. Cross-β polymerization and hydrogel formation by low-complexity sequence proteins. Methods 126, 3–11 (2017).

24. Cappannini, A., Forcelloni, S. & Giansanti, A. Evolutionary pressures and codon bias in low complexity regions of plasmodia. Genetica 149, 217–237 (2021).

25. Weber, L. M. et al. Essential guidelines for computational method benchmarking. Genome Biol. 20, 125 (2019).

26. Mier, P. et al. Disentangling the complexity of low complexity proteins. Brief. Bioinform. 21, 458–472 (2020).

27. Lim, K. G., Kwoh, C. K., Hsu, L. Y. & Wirawan, A. Review of tandem repeat search tools: a systematic approach to evaluating algorithmic performance. Brief. Bioinform. 14, 67–81 (2013).

28. Jarnot, P. et al. PlaToLoCo: the first web meta-server for visualization and annotation of low complexity regions in proteins. Nucleic Acids Res. 48, W77–W84 (2020).

29. Ziemska-Legiecka, J. et al. LCRAnnotationsDB: a database of low complexity regions functional and structural annotations. BMC Genomics 25, 1251 (2024).

30. Wootton, J. C. & Federhen, S. Statistics of local complexity in amino acid sequences and sequence databases. Comput. Chem. 17, 149–163 (1993).

31. Harrison, P. M. fLPS: Fast discovery of compositional biases for the protein universe. BMC Bioinformatics 18, 476 (2017).

32. Harrison, P. M. fLPS 2.0: rapid annotation of compositionally-biased regions in biological sequences. PeerJ 9, e12363 (2021).

33. Silva, J. M., Qi, W., Pinho, A. J. & Pratas, D. AlcoR: alignment-free simulation, mapping, and visualization of low-complexity regions in biological data. Gigascience 12, (2022).

34. Persi, E., Wolf, Y. I., Karamycheva, S., Makarova, K. S. & Koonin, E. V. Compensatory relationship between low-complexity regions and gene paralogy in the evolution of prokaryotes. Proc. Natl. Acad. Sci. U. S. A. 120, e2300154120 (2023).

35. Jorda, J. & Kajava, A. V. T-REKS: identification of Tandem REpeats in sequences with a K-meanS based algorithm. Bioinformatics 25, 2632–2638 (2009).

36. Newman, A. M. & Cooper, J. B. XSTREAM: A practical algorithm for identification and architecture modeling of tandem repeats in protein sequences. BMC Bioinformatics 8, 382 (2007).

37. R Core Team. R: A Language and Environment for Statistical Computing. (2024).

38. Wickham, H. et al. Welcome to the Tidyverse. J. Open Source Softw. 4, 1686 (2019).

39. Wickham, H. Ggplot2. (Springer International Publishing, Cham, 2016). doi:10.1007/978-3-319-24277-4.

40. Simon Garnier, N. R. R. R. A. P. C. M. S. and C. S. viridis(Lite) - Colorblind-Friendly Color Maps for R. (2024).

41. Wickham, H. Reshaping Data with the reshape Package. J. Stat. Softw. 21, (2007).

42. Hadley Wickham and Romain François and Lionel Henry and Kirill Müller and Davis Vaughan. dplyr: A Grammar of Data Manipulation. (2023).

43. Hadley Wickham and Davis Vaughan and Maximilian Girlich. tidyr: Tidy Messy Data. (2024).

44. Quinlan, A. R. & Hall, I. M. BEDTools: a flexible suite of utilities for comparing genomic features. Bioinformatics 26, 841–2 (2010).

45. Coombes, K. R., Brock, G., Abrams, Z. B. & Abruzzo, L. V. Polychromell: Creating and Assessing Qualitative Palettes with Many Colors. J. Stat. Softw. 90, (2019).

46. Thomas Lin Pedersen. patchwork: The Composer of Plots. (2024).

47. Taiyun Wei and Viliam Simko. R package ‘corrplot’: Visualization of a Correlation Matrix. (2024).

48. Sickmeier, M. et al. DisProt: the Database of Disordered Proteins. Nucleic Acids Res. 35, D786–D793 (2007).

49. Uversky, V. N., Oldfield, C. J. & Dunker, A. K. Showing your ID: intrinsic disorder as an ID for recognition, regulation and cell signaling. Journal of Molecular Recognition 18, 343–384 (2005).

50. Marcotte, E. M., Pellegrini, M., Yeates, T. O. & Eisenberg, D. A census of protein repeats. J. Mol. Biol. 293, 151–160 (1999).

51. Albà, M. M. & Guigó, R. Comparative Analysis of Amino Acid Repeats in Rodents and Humans. Genome Res. 14, 549–554 (2004).

52. Claus O. Wilke. cowplot: Streamlined Plot Theme and Plot Annotations for ‘ggplot2’. (2024).

53. Hadley Wickham and Jim Hester and Jennifer Bryan. readr: Read Rectangular Text Data. (2024).

54. Coletta, A. et al. Low-complexity regions within protein sequences have position-dependent roles. BMC Syst. Biol. 4, 43 (2010).

55. Mier, P. & Andrade-Navarro, M. A. The Conservation of Low Complexity Regions in Bacterial Proteins Depends on the Pathogenicity of the Strain and Subcellular Location of the Protein. Genes (Basel). 12, 451 (2021).

56. Gonçalves-Kulik, M. et al. Low Complexity Induces Structure in Protein Regions Predicted as Intrinsically Disordered. Biomolecules 12, 1098 (2022).

57. Dickson, Z. W. & Golding, G. B. Low Complexity Regions in Mammalian Proteins are Associated with Low Protein Abundance and High Transcript Abundance. Mol. Biol. Evol. 39, (2022).

58. Harrison, P. M. Patterny: A Troupe of Decipherment Helpers for Intrinsic Disorder, Low Complexity and Compositional Bias in Proteins. Biomolecules 15, 1332 (2025).

59. Kumari, B., Kumar, R. & Kumar, M. Low complexity and disordered regions of proteins have different structural and amino acid preferences. Mol. Biosyst. 11, 585–594 (2015).

60. Enright, J. M., Dickson, Z. W. & Golding, G. B. Low Complexity Regions in Proteins and DNA are Poorly Correlated. Mol. Biol. Evol. 40, (2023).

61. Uversky, V. N., Gillespie, J. R. & Fink, A. L. Why are ‘natively unfolded’ proteins unstructured under physiologic conditions? Proteins 41, 415–27 (2000).

62. Schneider, R. et al. Towards a robust description of intrinsic protein disorder using nuclear magnetic resonance spectroscopy. Mol. BioSyst. 8, 58–68 (2012).

63. Pavlović-Lažetić, G. M. et al. Bioinformatics analysis of disordered proteins in prokaryotes. BMC Bioinformatics 12, 66 (2011).

64. Necci, M. et al. Critical assessment of protein intrinsic disorder prediction. Nat. Methods 18, 472–481 (2021).

65. Das, S. et al. Sequence Complexity of Amyloidogenic Regions in Intrinsically Disordered Human Proteins. PLoS One 9, e89781 (2014).

66. Ntountoumi, C. et al. Low complexity regions in the proteins of prokaryotes perform important functional roles and are highly conserved. Nucleic Acids Res. 47, 9998–10009 (2019).

