## Supplementary Figures for "A Benchmarking Framework for Comparative Evaluation of Low-Complexity Region Detection Tools in the Human Proteome"

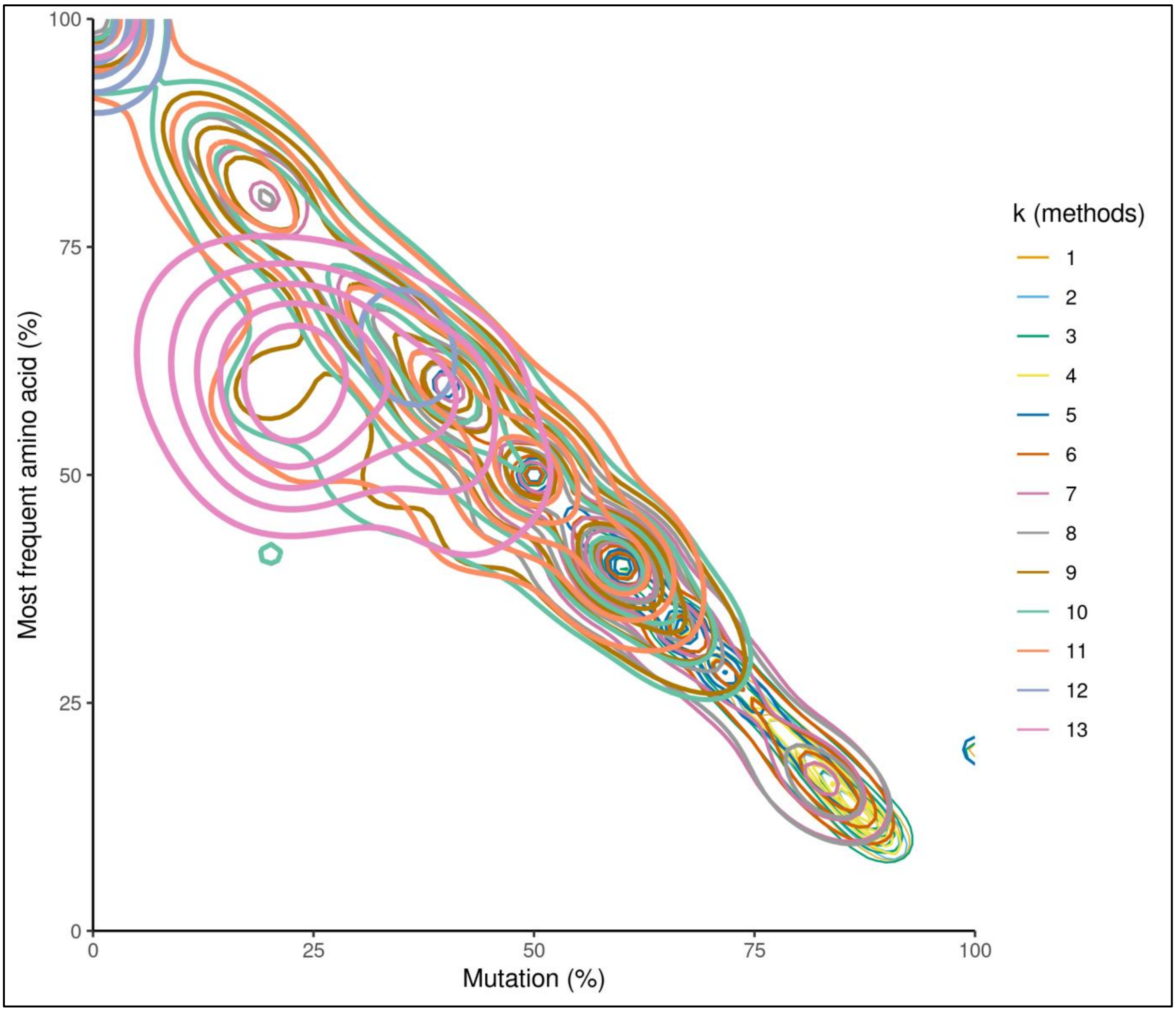


**Supplementary Figure 1. Shared LC manifold sharpened by consensus.** Kernel density contours of LC regions in mutation–composition space, colored by consensus tier (k). Contours indicate regions of increasing density.

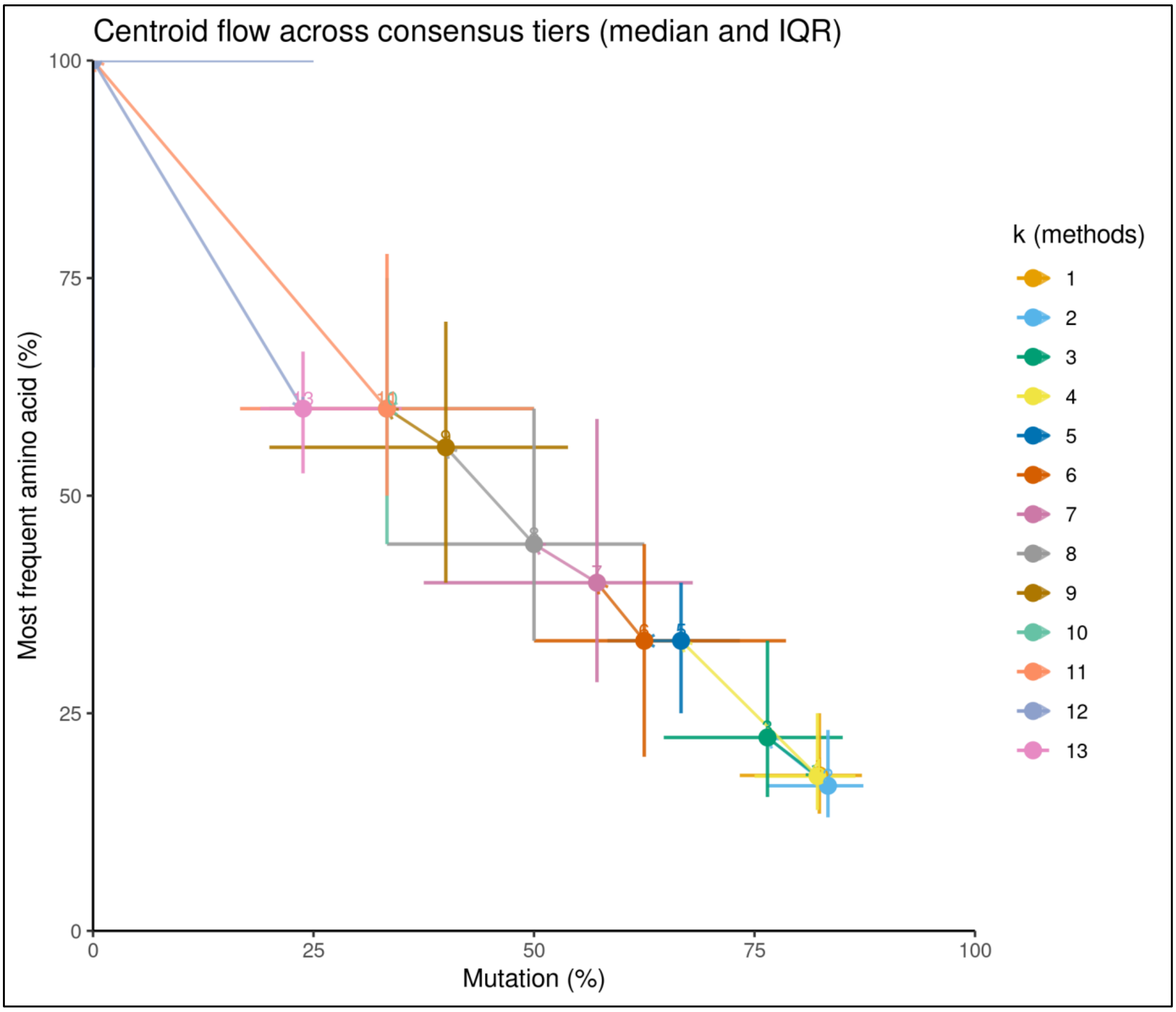


**Supplementary Figure 2**. **Consensus progression follows a directional complexity gradient.** Centroid flow plot showing median positions and interquartile ranges of mutation percentage and most frequent amino acid fraction for each consensus tier (k).

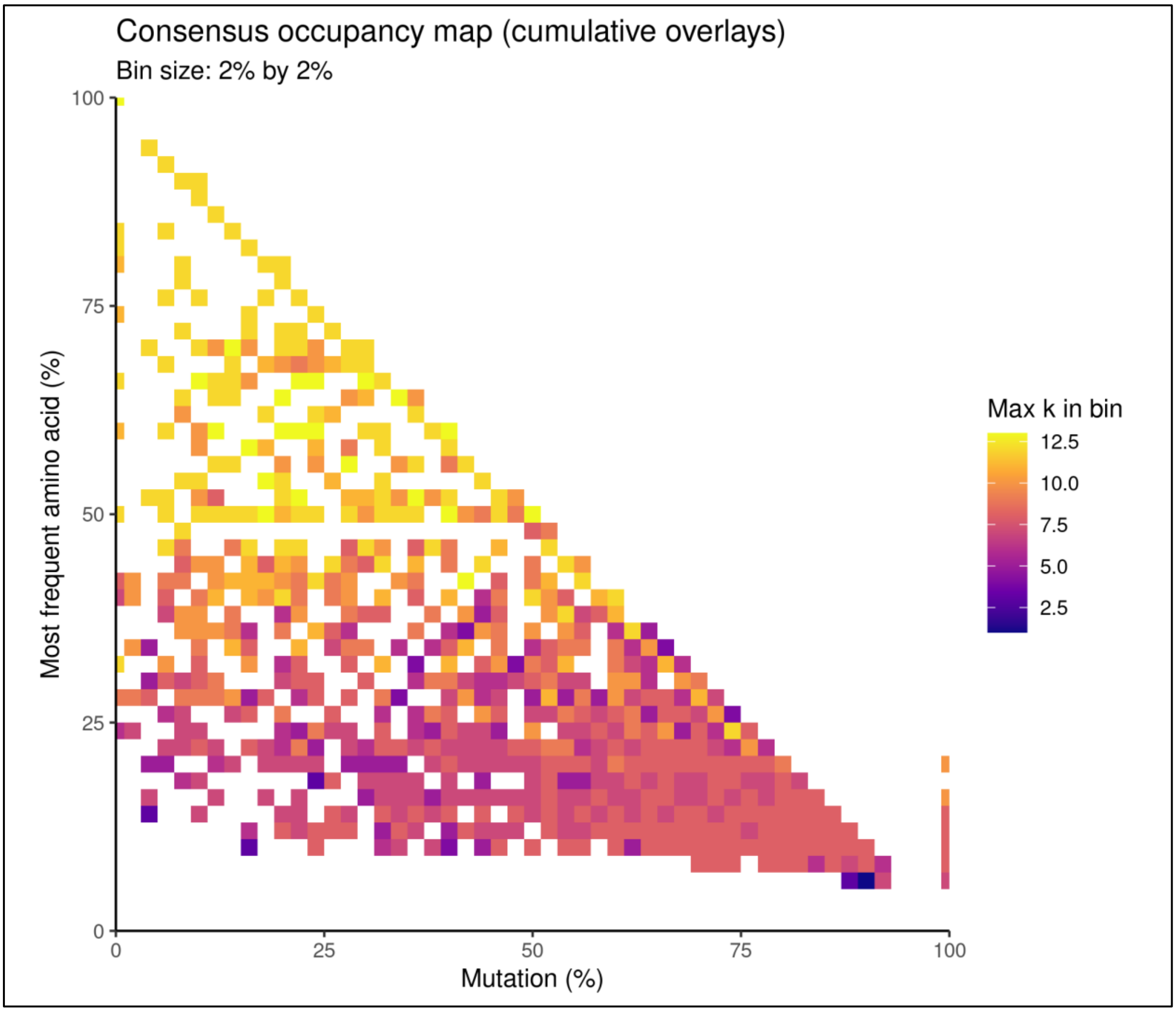


**Supplementary Figure 3. Maximal method agreement concentrates in the intermediate LC space.** Consensus occupancy heatmap showing the maximum number of overlapping methods (k) per 2% × 2% bin in LC space.

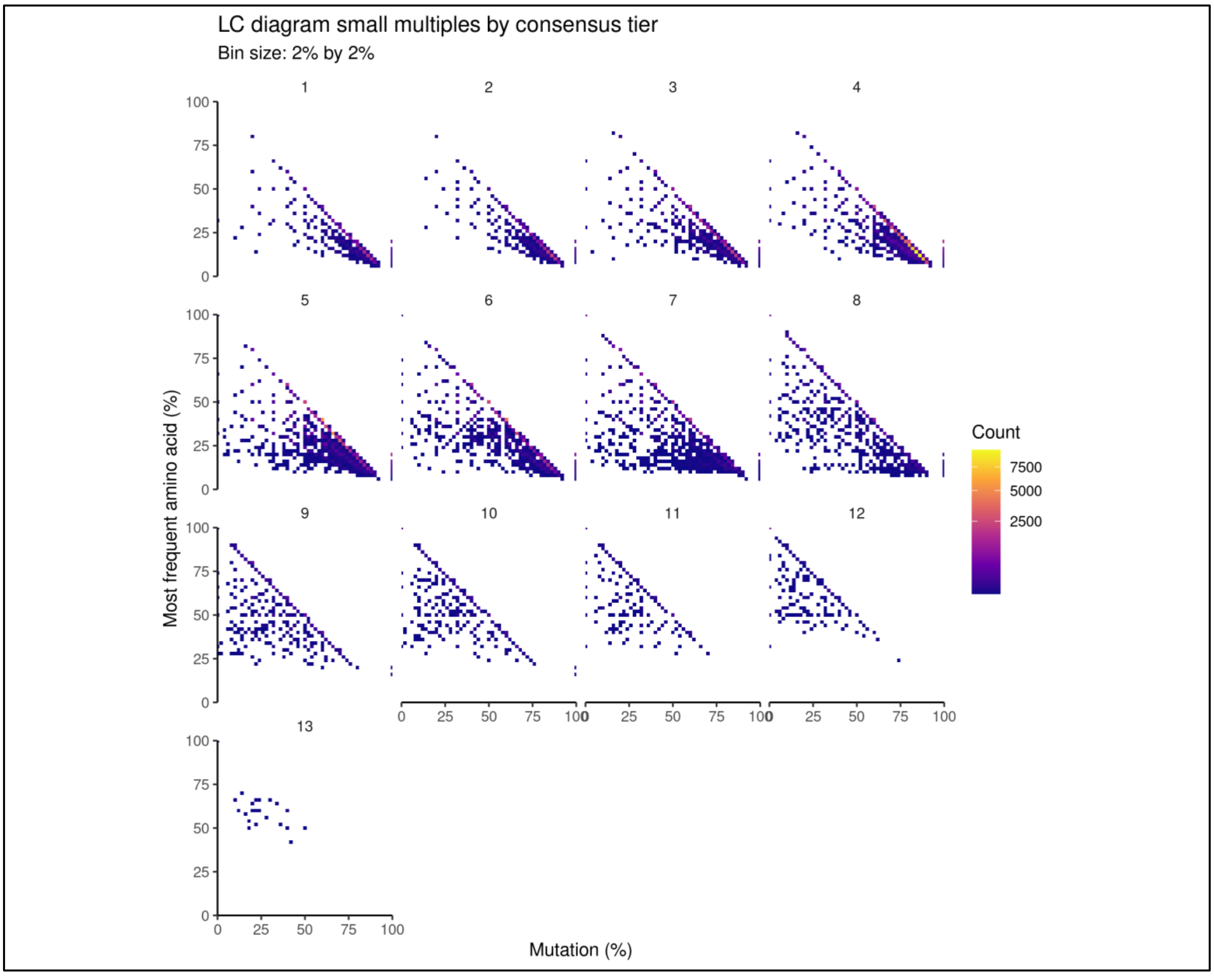


**Supplementary Figure 4. Increasing consensus selects for mutation-tolerant LC regions.** LC diagram small multiples stratified by consensus tier (k = number of methods). Each panel shows binned occupancy (2% × 2%) of regions in mutation percentage versus the most frequent amino acid fraction. Colour denotes bin counts.

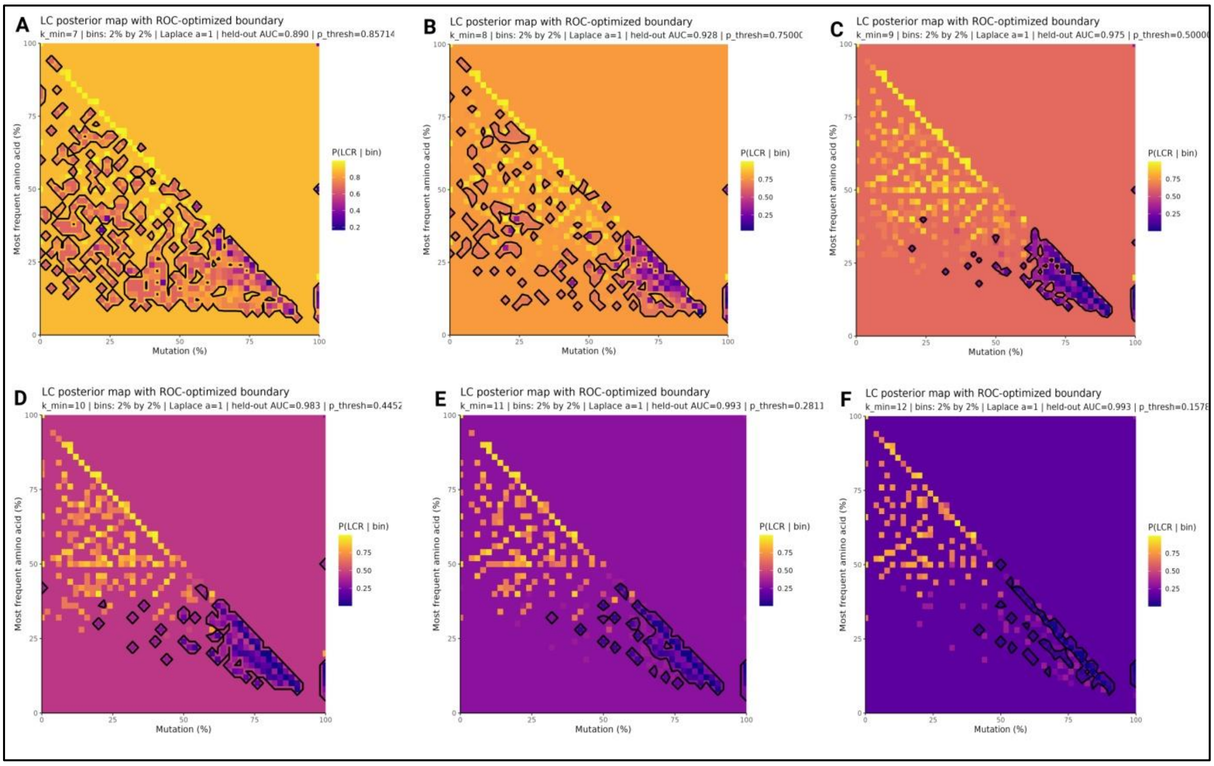


**Supplementary Figure 5. UniProt-derived LC posterior maps and ROC-optimised boundaries.** (A–F) LC posterior probability maps inferred from UniProt low-complexity annotations for increasing minimum window sizes (k = 7–12). Each panel displays P(LC | bin) across mutation percentage (x-axis) and most frequent amino acid percentage (y-axis), with warmer colours indicating higher LC probability. Black contours indicate ROC-optimised decision boundaries obtained from held-out validation data.

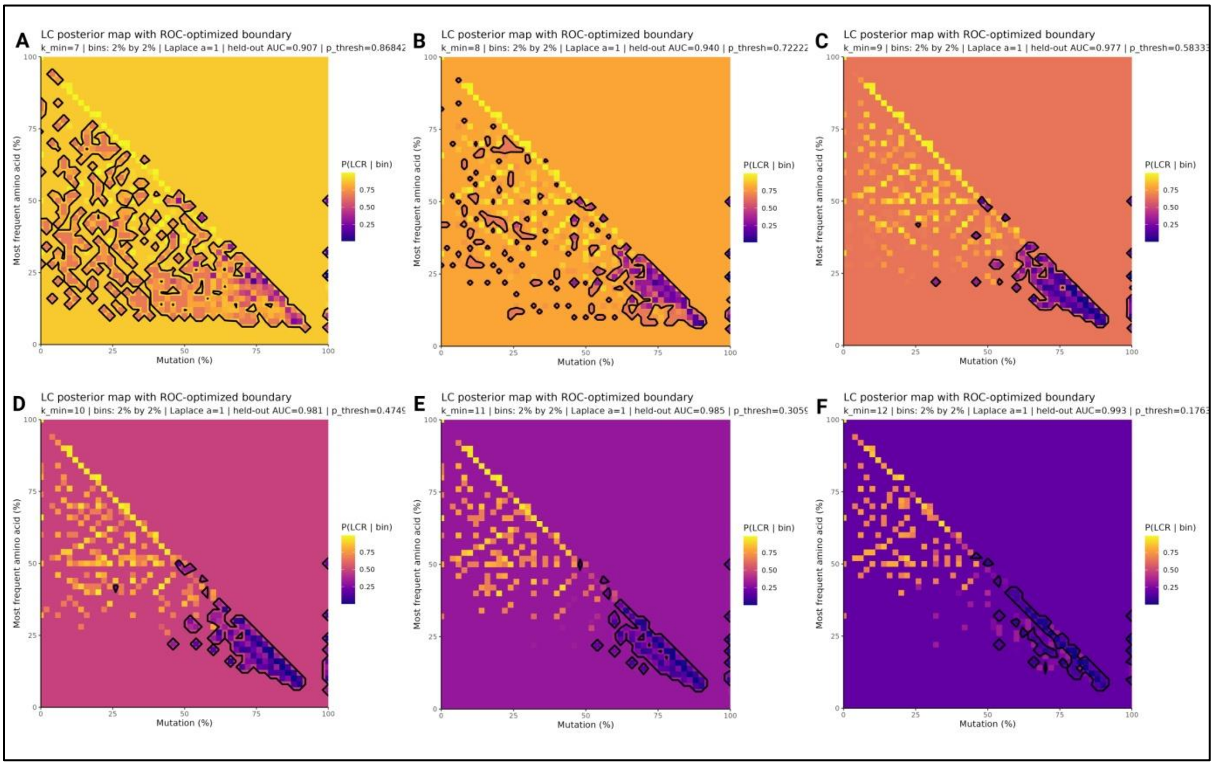


**Supplementary Figure 6. DisProt-derived LC posterior maps and ROC-optimised boundaries.** (A–F) LC posterior probability landscapes derived from DisProt-annotated intrinsically disordered regions across k = 7–12. Heatmaps show P(LC | bin) as a function of mutation percentage and compositional dominance. ROC-optimised boundaries (black contours) are broader and more fragmented at lower k, reflecting higher entropy tolerance in disorder annotations.

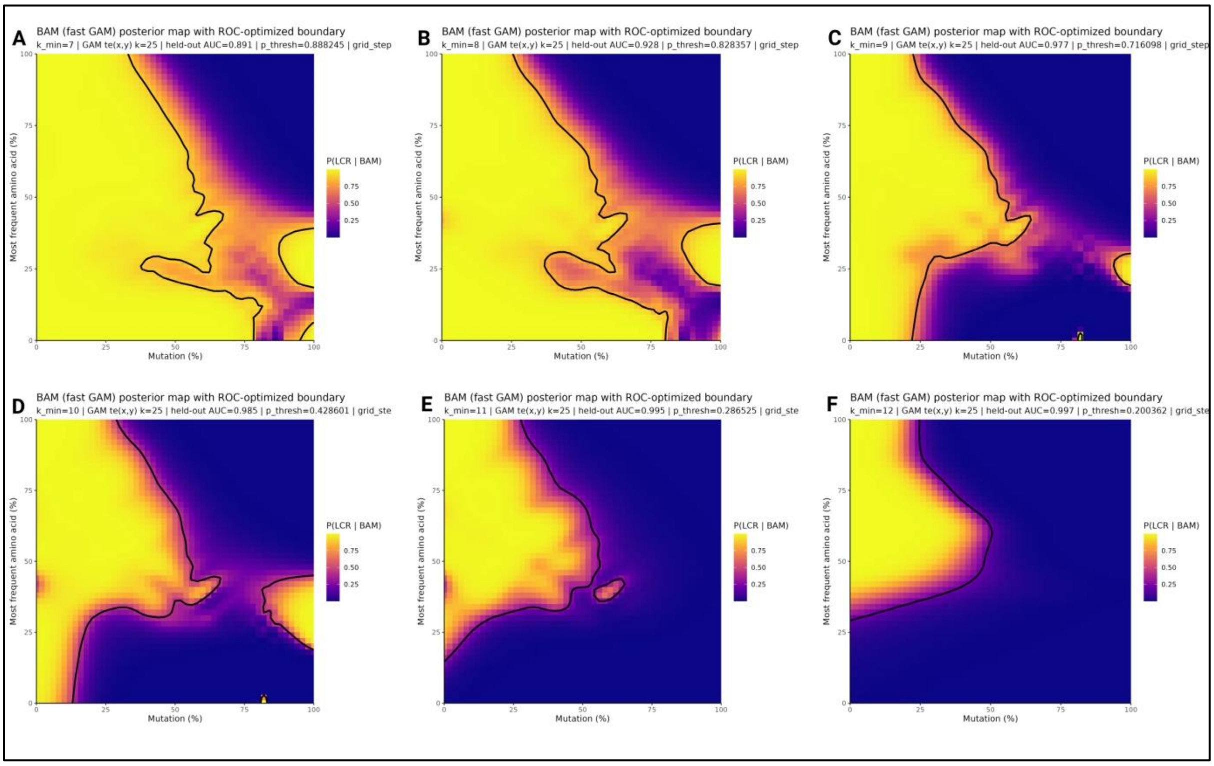


**Supplementary Figure 7. BAM LC posterior maps using UniProt annotations.** (A–F) Bayesian Additive Model (BAM) posterior probability surfaces trained on UniProt LC annotations for k = 7–12. Smooth posterior gradients depict P(LC|BAM) across mutation and compositional axes. Black curves indicate ROC-optimised decision boundaries derived from the continuous posterior surface.

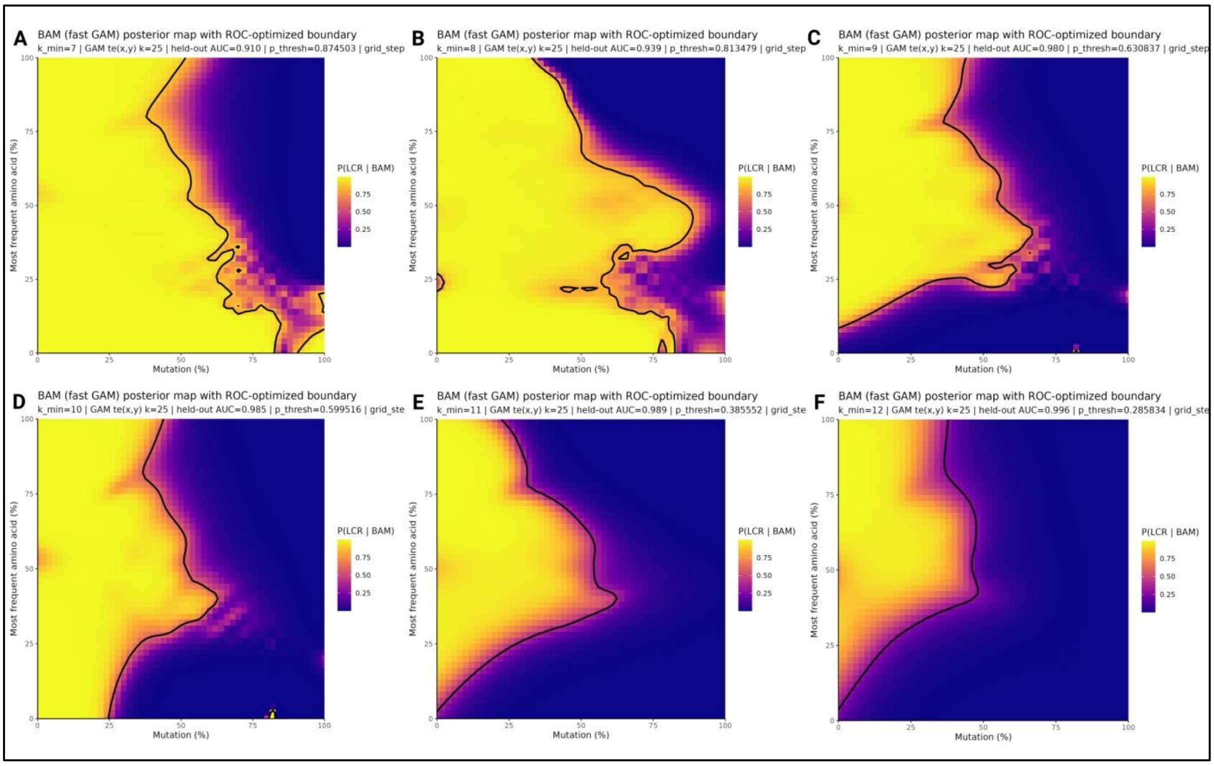


**Supplementary Figure 8. BAM (fast GAM) LC posterior maps using DisProt annotations.** (A–F) BAM-derived LC posterior probability maps trained on DisProt annotations across increasing k (7–12). Posterior surfaces reveal expanded LC space at lower k, with smooth entropy-driven transitions across mutation and compositional dimensions. ROC-optimised boundaries (black curves) remain broader than UniProt-based counterparts but contract and regularise with increasing k.

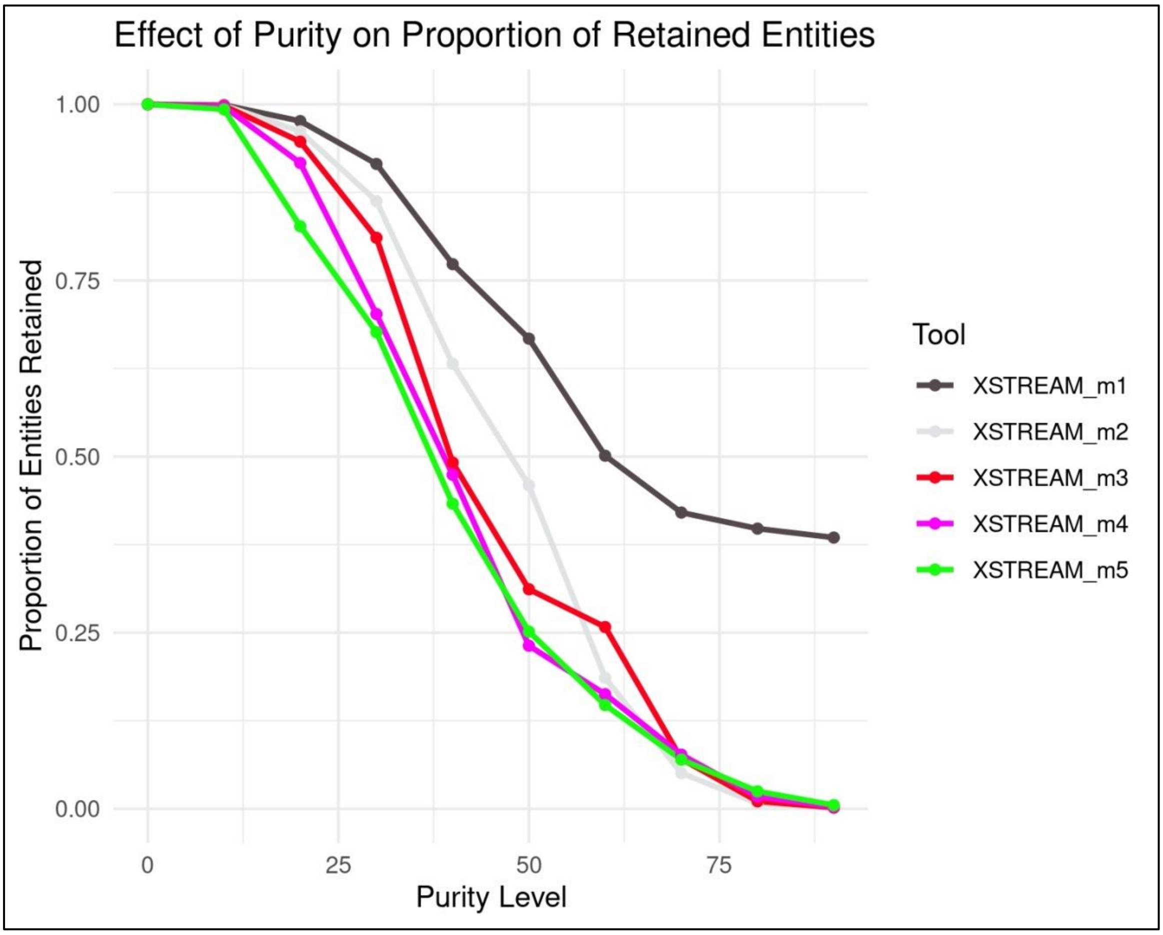


**Supplementary Figure 9. Retention of LCR purity by XSTREAM (at various -m parameters).** This figure shows the variation in LCR purity by XSTREAM (at various -m parameters). The x-axis represents the purity level, defined as the percentage of the most dominant amino acid within each LCR, while the y-axis denotes the proportion of total LCRs retained at or above each purity threshold. Each line corresponds to a different -m value, as indicated by the colour-coded legend. Steeper declines indicate detection of fewer highly pure (compositionally biased) regions, whereas gradual slopes reflect identification of more homogeneous LCRs. This comparison emphasisesXSTREAM varies in its sensitivity to amino acid repetitiveness, at various -m values.
